# Nonlinear Impacts of Herbivory on Plants Explain the Herbivory Paradox

**DOI:** 10.64898/2026.02.24.707788

**Authors:** Vincent S. Pan, Jared Adam, Daniel N. Anstett, A. Nalleli Carvajal Acosta, Tatiana Cornelissen, Andrea Galmán, Piper Haslup, Julia Karp, Xosé López-Goldar, Sylvie Martin-Eberhardt, Kelsen Ritter, Hellen Daiany Santos Lopes, Nicole E. Wonderlin, William C. Wetzel

**Affiliations:** Department of Integrative Biology, Michigan State University, East Lansing, Michigan, USA; W. K. Kellogg Biological Station, Michigan State University, Hickory Corners, Michigan, USA; Ecology, Evolution, and Behavior Program, Michigan State University, East Lansing, Michigan, USA; Land Resources and Environmental Sciences, Montana State University, Bozeman, Montana, USA; School of Integrative Plant Science, Section of Biology and the L. H. Bailey Hortorium, Cornell University, Ithaca, New York, USA; Department of Entomology, Michigan State University, East Lansing, Michigan, USA; Department of Plant Biology, Michigan State University, East Lansing, Michigan, USA; Center for Ecological Synthesis and Conservation, Universidade Federal de Minas Gerais, Belo Horizonte, Minas Gerais, Brazil; Department of Statistics, Williams College, Williamstown, Massachusetts, USA; School of Biological Sciences, Illinois State University, Normal, Illinois, USA; Departamento de Biodiversidade, Universidade Estadual Paulista Júlio de Mesquita Filho, Rio Claro, São Paulo, Brazil; Department of Integrative Biology, The University of Texas at Austin, Austin, Texas, USA

**Keywords:** Herbivory, Interaction strength, Meta-analysis, Nonlinear averaging, Plant fitness, Plant-herbivore interactions, Tolerance

## Abstract

Herbivores strongly shape plant ecology and evolution. Yet plants typically sustain low, tolerable levels of damage, raising the question of how herbivory influences plant biology despite often weak fitness effects. Analyzing 1,145 datasets of the effects of damage on fitness for 103 plant species worldwide, we show that nonlinear tolerance to damage is one explanation for this paradoxical contradiction. Plants are tolerant of low levels of damage, but infrequent, severe damage levels disproportionately reduce plant fitness. Models incorporating nonlinear tolerance suggest that disproportionate tolerance to low levels of damage stabilizes population dynamics and promotes coevolutionary feedback with herbivores. Pervasive nonlinear tolerance shows consistent patterns across environmental gradients, may alter the stability of food webs, and may explain why herbivory matters despite the world being green.

## INTRODUCTION

The extent to which herbivores influence plant fitness has long been a source of controversy (*1*–*3*). Herbivory has well-documented consequences for plant community composition and has driven the repeated evolution of a remarkable diversity of plant defense traits (*4, 5*). Yet empirical estimates of herbivore effects on plant fitness are often minimal under typical field conditions (*6, 7*), with many plants displaying strong tolerance or even overcompensation to damage (*8*). A paradoxical contradiction thus exists between the major consequences of herbivory at larger scales and its minor effects on individual plant fitness. Understanding how these minor fitness impacts scale up to shape plant ecology and evolution remains an important unsolved challenge (*9, 10*). Solutions to this herbivory paradox have emphasized evolved plant defenses that mitigate the effects of herbivory in modern day (*11*), but fossil evidence indicates that the effects of modern-day herbivory are if anything stronger (*12*). Other solutions have emphasized that through other interactions, such as competition, herbivory can indirectly increase its importance (*13*), but the prevalence of these higher-order interactions remains an open problem. Here, we propose and test an alternative hypothesis that nonlinear effects of herbivore damage on plant fitness could sustain the importance of herbivory, even when the fitness effects are typically negligible.

The relationship between herbivore damage and the resulting reduction in plant fitness is an integral component of all models involving plant–herbivore interactions and plant demography. This relationship, which we call the *damage function*, therefore has broad applications in predicting population, community, and crop yield dynamics (*14, 15*). The function is hypothesized to display a variety of shapes (Figure 1A; (*16*)), ranging from disproportionately weak to strong tolerance to low relative to high levels of damage (or *sensitive* versus *insensitive* for short). Yet despite repeated calls to characterize the damage function (*17, 18*), existing analyses are usually limited to one species (*19*) and often assume a linear relationship (*20*). This mismatch may be significant. Although little theory exists for nonlinear damage functions (*21*), work on other functions central to demography, such as consumer functional responses (*22*), shows that nonlinearity can qualitatively alter dynamics, sometimes even reversing classical intuitions (*23, 24*). Likewise, empiricists have long recognized that tolerance to herbivory can be as important as the resistance traits that prevent herbivory (*25, 26*), but idiosyncratic experimental methodologies and definitions of tolerance have impeded quantitative syntheses of broad patterns (*17, 20, 25*). Understanding how the damage function can vary across biological contexts and its implications for dynamics is therefore a crucial gap.

**Figure 1.**
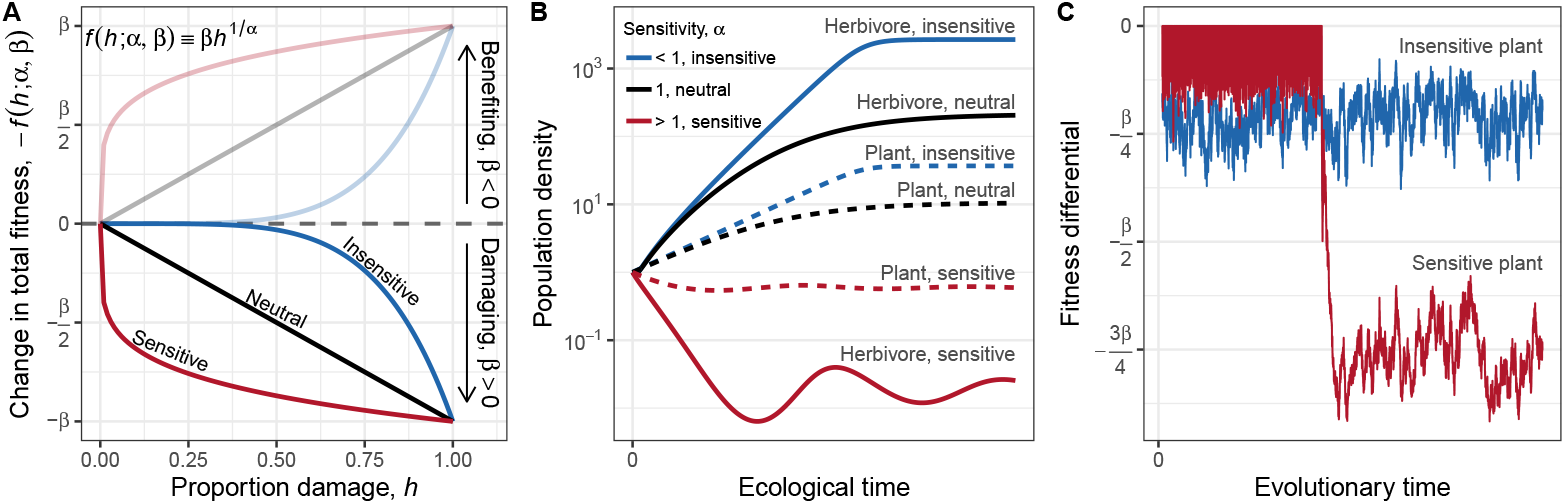
Impact of damage function on the population and coevolutionary dynamics of plants and herbivores. (A) The degree to which a given proportion of tissue damage *h* reduces plant fitness, or the *damage function f* (*h*), can take on many shapes well described by a family of power functions. These functions are defined by the damage intensity parameter *β* (reduction in fitness at total damage, *h* = 1) and damage sensitivity parameter *α*. When *β* > 0, tissue damage reduces plant fitness (opaque lines), whereas when *β* < 0, damage increases plant fitness (transparent lines). When *α* = 1, the plant is equally sensitive to low and high levels of damage; when *α* > 1 or *α* < 1, the plant is disproportionately more sensitive to low and high levels of damage, respectively. (B) Over ecological time, greater insensitivity allows the plant and herbivore population to achieve higher equilibrium density and stability (Appendix 2, Table S5, Figure S2-S3). Over evolutionary time, insensitivity, in contrast to sensitivity, leads to the evolution of one, rather than two, equilibrium mean damage level, making highly detrimental levels of damage less frequent (Appendix 3, Figure S4-S5). The results of (B) and (C) together provide conditions ripe for a mismatch between the minor fitness impacts of herbivory and its major ecological and evolutionary consequences.

The importance of damage functions in plant-herbivore biology led us to hypothesize that it could also explain the herbivory paradox. Observations that plants often exhibit strong regrowth in response to minor damage (*8, 18*) and the prediction that damage releases less productive leaves from self-shading in canopy models (*15*) suggest that plants should disproportionately tolerate low levels of damage, displaying insensitivity. We hypothesize that if insensitivity is indeed widespread, the steep change in fitness at high damage levels would cause instances of high damage, even if rare, to disproportionally contribute to the overall effects on plant fitness. Simply stated, insensitivity could amplify the importance of infrequent but severe damage through nonlinear averaging, offering one overlooked solution to the herbivory paradox. Our hypothesis accords with long standing notion that the effects of herbivory are driven by a few bursts of top-down control, such as during an herbivore outbreak, between long periods of stasis (*6, 27, 28*). Concordantly, recent evidence indicates that low-frequency, high damage levels account for most of the herbivore damage in nature even though plants on average receive low damage (*29, 30*).

Using nonlinear models, we reanalyzed 1,145 datasets of damage-fitness relationships collected from 100 studies, spanning 103 species and 116° of latitude (Appendices 1 and 4, Table S1-S4, Figure S1). We included studies that measured plant fitness components across experimentally damaged plants with at least four unique damage levels. We focused on studies that randomly assigned artificial damage because real herbivores may adjust their feeding rate in ways linked to plant fitness, thus introducing bias (*31, 32*). We approximated damage functions with a family of power functions *f* (*h*; *α, β*) ≡ *βh*^1/*α*^, where *h* is the proportion of standing biomass removed, *β* scales the intensity of reduction in plant fitness, and *α* controls the sensitivity to low levels of damage (Figure 1A). Difficulty in quantifying lifetime plant fitness led us to decomposed overall fitness into fitness components across different episodes of selection (*33*). We thus modeled the damage function for individual fitness components 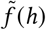, on average informing the shape of *f* (*h*) and the minimum magnitude of its effect (Appendix 1). Here, we first explored how including nonlinear damage functions in consumer-resource (*21*) and coevolutionary (*34*) models affect dynamics. We then report on the shape of damage functions and how nonlinearity magnifies the mean and variance of damage effects on fitness. Finally, we tested longstanding hypotheses about the variation in tolerance to herbivore damage across environmental gradients, plant functional groups, and life history.

## RESULTS AND DISCUSSION

### Damage function nonlinearity can control dynamics

We found that the shape of damage functions can qualitatively affect model dynamics. Over ecological time, insensitivity, in contrast to sensitivity, may promote herbivore regulation of plant populations (Figure 1B, Appendix 2). It also may increase the stability and equilibrium density of both plants and herbivores, especially if damage reduces plant fitness through lowering plant fecundity instead of survival. This result provides further evidence that insensitivity should be widespread. Over evolutionary time, insensitivity may lead to a single coevolutionary regime where moderately defended plants sustain damage they can tolerate (Figure 1C, Appendix 3). The high tolerance predicted is consistent with empirical evidence (*6, 7, 29*) and importantly, may provide a necessary condition for the herbivory paradox. Conversely, sensitivity may weaken coevolutionary feedback, often leading to bistable regimes where plants sustain either extremely low or high damage levels, a pattern rarely seen in terrestrial ecosystems (*29*). These results illustrate that nonlinear damage functions are important for the ecological and evolutionary dynamics of plant-herbivore interactions from which the herbivory paradox may readily arise.

### Damage functions often display insensitivity

We found that plant fitness was on average strongly insensitive to low levels of damage, declining slowly at low levels of damage, but rapidly at high levels of damage (Figure 2A, S6-S8). Indeed, a typical field-average of 11% damage (Table S3) reduced plant fitness component on average by a small 1.7% [1.4%, 1.9%] (mean [95% confidence interval]), whereas 95% damage caused a sizable 27% [25%, 30%] reduction. This ∼ 16 times greater reduction in fitness comes from only a ∼ 8.6 times increase in damage, consistent with our hypothesis that high damage levels have a disproportionately large influence on plant fitness.

**Figure 2.**
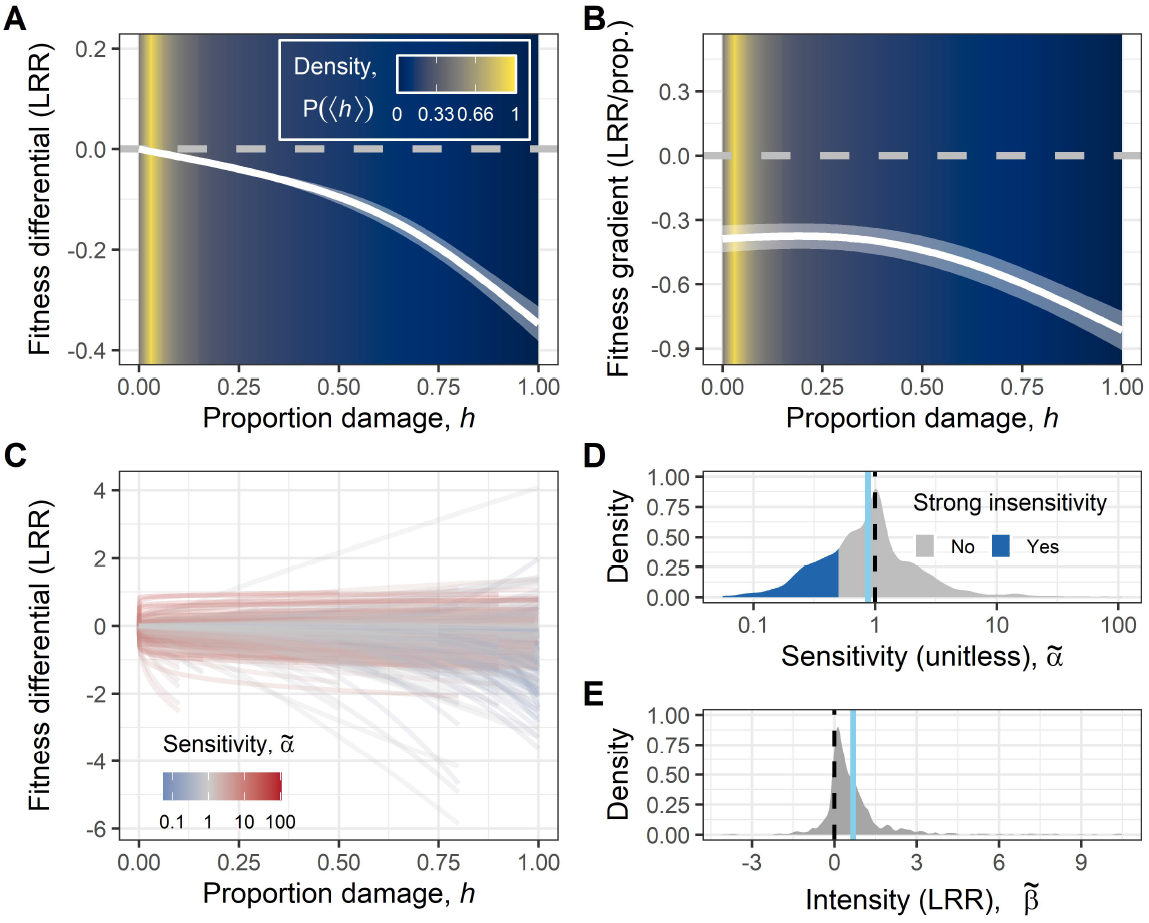
Patterns of damage function 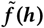 in an episode of selection. (A) The average fitness differential 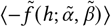 and (B) fitness gradient 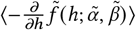 in an episode of selection across all datasets show that plants in general can tolerate low, but not high, levels of damage. Lines and ribbons show mean and 95% confidence intervals. LRR stands for log response ratio, where the undamaged condition (*h* = 0) is used as a reference. Typical levels of mean herbivore damage (Table S3), shown by background color, tend to fall in a region where plants are insensitive to damage. (C) The best-fit negative damage functions (lines) in an episode of selection 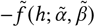 for each dataset show considerable variation. (D-E) Density plots show the distribution of maximum *a posteriori* estimates of fitted parameters. The black dashed lines show the null values representing a null damage function of a flat 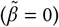 line 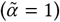. The blue solid lines show the mean parameter values. Parameters that were classified as strong insensitivity 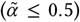 are shaded blue. More details are reported in Appendices 1, 4, and 5, Table S6.

The pattern of insensitivity is robust. Examining the slope between plant fitness and damage (fitness gradient), a more common measure of tolerance (*20*), we likewise found that tolerance declined by roughly twofold from low to high levels of damage, rather than being constant (Figure 2B). When we classified individual damage functions to explore their heterogeneity (Figure 2C-E), we found that 93% of datasets showed significant deviation from linearity, with insensitivity (61%) being roughly twice as common as sensitivity (32%). Moreover, only 13% of datasets showed a significantly positive effect of damage on fitness (negative effects: 84%), suggesting that damage seldom confers direct fitness benefits to plants (*3*). These results are robust to our assumed shape of the damage function, as supported by nonparametric analyses (Figure S7) and by evidence that sigmoidal and hormetic functional forms (*14, 35*) are relatively uncommon (Appendix 4, Figure S6).

These findings agree with our hypothesis that damage functions typically display insensitivity, with most plants able to disproportionally tolerate the low levels of damage they typically experience (*8, 15, 18*). Insensitivity could occur because plants have excess organs or tissues, such as shaded leaves or dormant meristems, or resources in reserve that can be mobilized for tissue repair (*15, 25*); although, these losses can only be compensated up to a point. According to models of tolerance based on limiting resources (Liebig’s law) (*36*), widespread insensitivity could also occur because herbivores generally do not reduce plant access to limiting resources, such as soil nutrients. Taking a broader perspective, the minimal fitness impact herbivores typically have on plants is consistent with existing work on food webs and suggests the weak effects of herbivores may not be unusual among other types of trophic interactions as is often believed (*1, 2*). Indeed, empirical estimates of trophic interaction strengths are typically weak (*37*), and this weakness may be prevalent because it promotes community stability (*23, 24*).

### Implications of insensitivity

We estimated the extent to which the disproportionate effect of high damage on plant fitness can inflate the average strength of damage effects. We found that this inflation was strongest for plants that were more insensitive to low levels of damage and notably, when the effect of average damage was weak. Thus, the inflation from high levels of damage effectively set a minimum magnitude on the effect of herbivory (Appendix 4). Simulations using field data from a global survey of herbivory patterns (*29*) confirmed that these high damage-levels which plants cannot tolerate are often enough to maintain significant effects of herbivore damage on plant fitness (Figure S9). Indeed, damage functions with the strongest insensitivity to low levels of damage (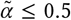, 24% of functions, Figure 2D) resulted in at least a 4.9-fold amplification in the fitness effects. The hypothesis that infrequent, high damage levels can sustain the importance of herbivory therefore lies in the phenomenon that nonlinear averaging prevents the effect of herbivore damage from becoming trivial, serving as one important solution to the herbivory paradox.

Because variation in interaction strengths can be as important as the mean strength (*29*), we also investigated the extent to which the disproportionate effect of high damage-level on plant fitness exacerbates variation in interaction strength. We found that in cases where the damage function exhibits strong insensitivity 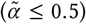, the most severe 10% of damage levels represent at least 58% of the total leaf area damaged, but our analysis showed that those damage levels explain at least 87% of the total reduction in fitness from herbivory. Such strong inequality in fitness impacts is important because when highly detrimental herbivory levels are infrequent, short-term studies likely miss these highly influential events. Herbivore damage may therefore appear to have trivial effects on plant fitness when the long-term consequences are not trivial. This dominance of plant-herbivore interaction by a few strongly influential events may also alter community dynamics. For example, theory suggests that when all but a few interactions are weak, it could stabilize food webs by dampening oscillations between strongly interacting plant–herbivore pairs (*24*). Likewise, under temporal fluctuations in damage, variation in interaction strengths combined with differences in the degree of nonlinear tolerance between species can potentially stabilize plant coexistence (*38*).

### Influence of environmental conditions

Understanding the context under which plants exhibit greater tolerance can explain patterns of plant biogeography and how plants respond to global change but a synthesis of tolerance to herbivory is lacking. Therefore, we examined how the shape of the damage function varied along major environmental gradients. The Latitudinal Herbivory-Defense Hypothesis, which posits that plant defense and herbivore damage increase towards the tropics (*29, 39*), attracted much attention for its role in explaining the latitudinal diversity gradient (*40*). An often-neglected component of this hypothesis is that tolerance to herbivore damage may also decrease with latitude (but see (*41*)). If so, greater damage may not translate to stronger interactions in the tropics. To test this hypothesis, we analyzed the parameters of the damage functions 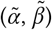 using data from field studies with georeferenced sites (*n* = 864). We split our analysis of damage sensitivity by whether damage increased or decreased plant fitness because fitness is maximized when plants are sensitive to beneficial effects and insensitive to detrimental effects of herbivory.

We found that plants at lower latitudes indeed suffered less reduction in reproduction due to damage, although these plants were less likely to survive (Figure 3A-C). Compared to temperate (±45°) plants, tropical (0°) plants were roughly as tolerant of the average level of damage (11% damage; growth: 2.1% [-1.2%, 5.2%]; reproduction: −6.0% [-12%, 0.10%]; survival: −0.52% [-3.7%, 2.5%]). However, at high levels of damage, tropical plants maintained moderately greater reproduction and growth, but at the expense of a substantial reduction in survival (95% damage; growth: 34% [11%, 51%]; reproduction: 44% [22%, 60%]; survival: −170% [-370%, −50%]). These results suggest that there are latitudinal gradients in tolerance, but they differ by life-history components, indicating that herbivory may exert fundamentally different selective pressures on plant strategies across latitudes (*42*). While we do not find definitive support for the Latitudinal Herbivory-Defense Hypothesis, we do find that plants which we expect are demographically more sensitive to changes in survival (e.g., slower growing plants), may have stronger interactions with herbivores in the tropics. Conversely, plants that rarely die from damage may respond more strongly to fluctuations in herbivore damage, which recent evidence indicates are more pronounced at higher latitudes (*29*).

**Figure 3.**
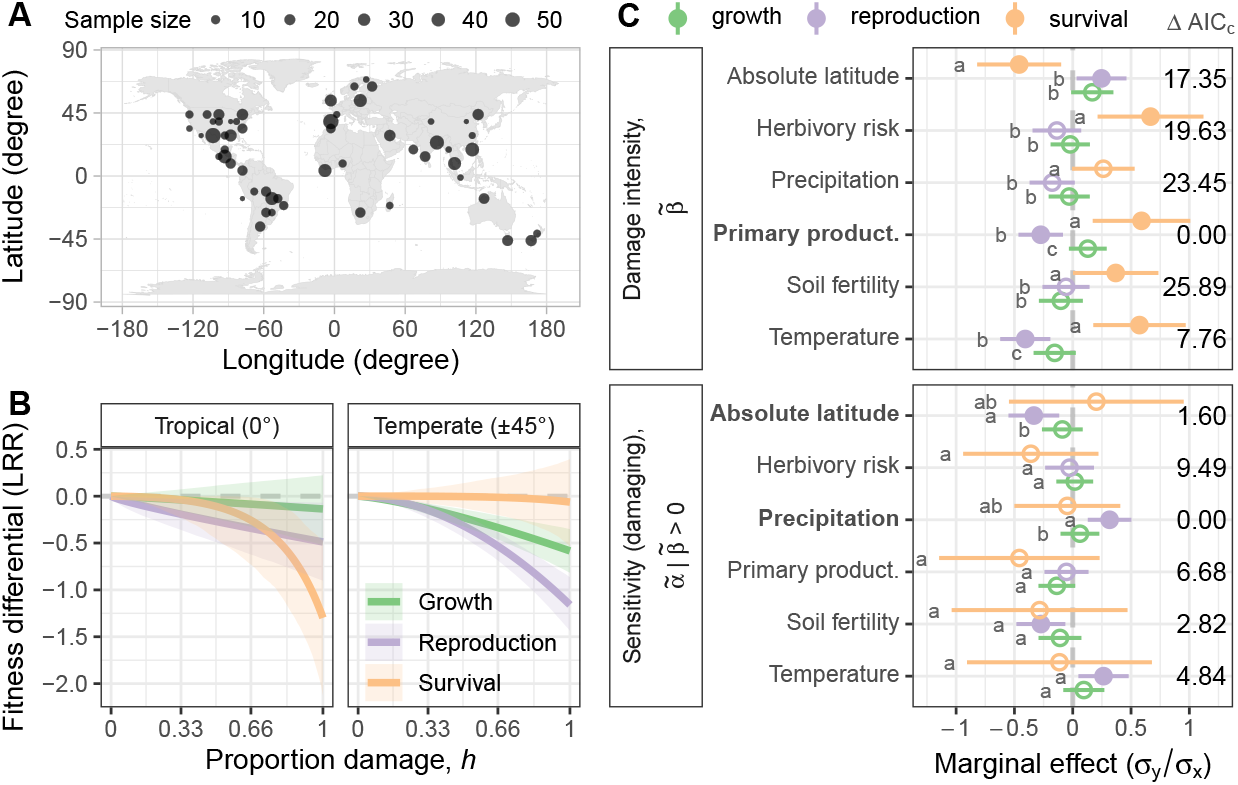
The environmental correlates of the damage function 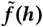. (A) Geographical distribution of sampling at 5° × 5° resolution. Point size is scaled to the number of samples in each grid cell. (B) The negative damage function 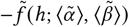 of each life-history component displays different shapes across latitude. Lines and ribbons show mean and 95% confidence intervals. LRR stands for log response ratio, where the undamaged condition (*h* = 0) is used as a reference. (C) The marginal effect of predictors on different damage function parameters. Points and ribbons show mean and 95% confidence intervals. Filled circles indicate significant parameters. Top predictor(s) based on ΔAIC_c_ are bolded. Different letters next to each effect indicate significant differences between marginal effects by life-history variable. More details are reported in Appendices 1 and 5, Table S7, Figure S10.

For a more mechanistic understanding of what might underlie the observed latitudinal gradient in tolerance, we explored five additional environmental variables potentially influencing plant performance (herbivory risk, precipitation, temperature, soil fertility, primary productivity, Figure 3C). Greater herbivory pressure is thought to select for increased tolerance to herbivore damage (*43*). Warmer, wetter, more productive environments with more nutrients are more amenable to plant growth and can either increase or decrease tolerance depending on the hypothesis (*35, 36, 44, 45*). We explored which variables best predicted damage function parameters using corrected Akaike Information Criterion (AIC_c_). We found that mean annual precipitation and absolute latitude best explained damage sensitivity 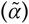, whereas net primary productivity best explained damage intensity 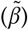 and both parameters overall (ΔAIC_c_ = 5.3, Figure 3C, S10). Lower sensitivity in more arid environments may stem from water being the main limiting resource, masking the effect of carbon loss due to herbivores (*36*). That tolerance in reproduction, but not survival, increased with productivity is consistent with the view that faster growing plants are more able to replace tissue lost from damage (*44, 45*). Greater competition in more productive environments, however, may limit the establishment of plants that experienced herbivory. These results suggest that the damage function is best explained by bottom-up factors and greater productivity toward the tropics may provide a tentative mechanism for the latitudinal gradient in tolerance.

### Influence of life history and functional groups

Finally, we tested long standing hypotheses about how tolerance of herbivore damage varies across plant life history and functional groups. In general, damage to reproductive organs and damage early in a plant’s development are thought to be more detrimental to fitness (*46, 47*). Annuals, graminoids, and herbaceous plants are considered more tolerant to damage than perennials and woody plants due to their ability to replace consumed tissue (*2, 26, 44, 45*). Domesticated plants are considered less tolerant to damage because traits important to tolerance, such as branching and tillering, are often artificially selected against (*48*). These assumptions underlie major evolutionary hypotheses of plant function, including those predicting allocation of herbivore defenses between organs (e.g., Optimal Defense Theory (*49*)) and between species (e.g., Resource Availability Hypothesis (*44*)). However, they have only been tested systematically in a few species and without considering nonlinearity (*26*).

We found surprisingly minor differences in tolerance across plants overall (Figure 4). Across all subgroups, plants remained largely insensitive to the negative effects of damage, neither sensitive nor insensitive to the positive effects of damage, and suffered a moderate reduction in fitness at total damage. Nevertheless, some differences did agree with existing hypotheses of tolerance. Plants were more insensitive to damage to stems or shoots than damage to leaves (−28% [-48%, −0.91%]) possibly because plants otherwise exhibited strong apical dominance and can reactivate suppressed buds in reserve (*25*). Damage during the fruiting stage reduced fitness more than any other stage (34% [6.7%, 70%]), possibly because photosynthates may be more limiting during the demanding process of seed filling. Damage intensity was strongest for reproduction, being 37% [17%, 59%] and 92% [32%, 179%] stronger than the intensity of growth and survival, respectively. Conversely, damage sensitivity of survival was lower compared to the damage sensitivity of reproduction (−42% [-65%, −4.3%]) and growth (−44% [-66%, −9.0%]). In other words, plant survival suffered the least at low levels of damage, whereas plant reproduction suffered the most at high levels of damage. These results are consistent with the view that herbivore damage primarily affects plant fitness through changes in fecundity because plants exhibit a strong ability to regrow and survive damage (*18*). As well, survival exhibited strong insensitivity because most plants survived, and the response’s upper-bound at no deaths results in saturation. Theory suggests that greater damage intensity on fecundity tends to destabilize plant-herbivore population dynamics, while lower sensitivity of survival has a stabilizing effect (Appendix 2). Together, plant responses to herbivore damage do not vary predictably across plant life-history strategies, domestication, or growth form. Rather, tolerance may be better explained by the plant’s environmental conditions and the organ and life-history component which damage affects.

**Figure 4.**
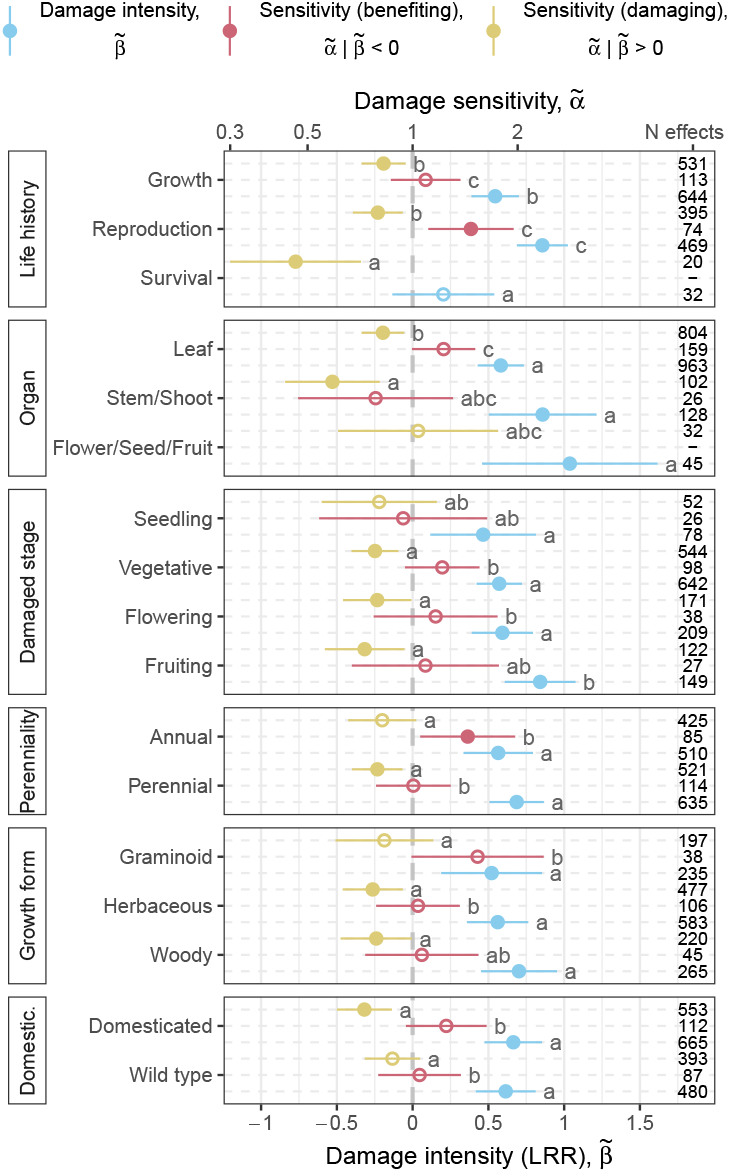
The average damage-function parameters across life history and functional groups. Points and ribbons show mean and 95% confidence intervals. LRR stands for log response ratio. Damage sensitivity is unitless. Filled circles indicate significant parameters. We displayed group means for when there are at least fifteen effect sizes from five different studies. Different letters next to each effect indicate significant differences between parameters (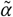 and 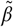). Aside from a few minor differences, plants remained largely insensitive to the negative effects of damage, neutrally sensitive to the positive effects of damage, and suffered a moderate reduction in fitness at total damage. There was a weak negative correlation between damage sensitivity 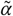 and damage intensity 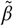 (*r* = −0.22 [-0.28, −0.16]). More details are reported in Appendices 1 and 5, Table S8.

## CONCLUSION

Pervasive nonlinear tolerance to damage offers one solution to the paradoxical contradiction between minor fitness impacts of herbivory and its major ecological and evolutionary consequences. Indeed, damage insensitivity makes high levels of damage disproportionately influential for overall fitness. While herbivore damage is low on average, its extreme variation makes high levels of damage surprisingly common. For example, fir and spruce herbivore damage vary by 10 – 80% in a decade (*50*) and many herbivorous insects fluctuate in density by more than two orders of magnitude (*28, 51*). Together, our findings support the view that plant-herbivore interactions may be episodic, with long periods of commensalism interspersed with bursts of consumer control. They add to a growing body of evidence that the pace at which the consequences of species interactions emerge in ecological and evolutionary signals is not gradual (*28, 52*), potentially limiting the efficiency of natural selection (*53*) and the predictability of ecological dynamics.

## ACKNOWLEDGMENTS

We thank Gideon Bradburd, Jeff Conner, Chris Klausmeier, Rick Karban, Kadeem Gilbert, Lauren Sullivan, Matt Forister, and Eric LoPresti for helpful discussion. Michael Kalwajtys helped us collect the preliminary data. This is Kellogg Biological Station Contribution no. ??.

## Author contributions

VSP and WCW conceived of the study and led the data collection. All authors contributed to the study design and collected the data. VSP analyzed the theoretical models and data, and wrote the manuscript. All authors provided editorial support and approved the manuscript. Middle authors are listed in alphabetical order.

## Data and materials availability

All code and data are deposited at Dryad (??). A private for peer-review link to the materials is here: ??

## SUPPLEMENTARY MATERIALS

Materials and methods (Appendix 1)

Supplementary text (Appendices 2 to 5)

Figure S1 to S10

Table S1 to S8

