## Supplementary materials for "Nonlinear Impacts of Herbivory on Plants Explain the Herbivory Paradox"

February 24, 2026

**This PDF file includes:**

Materials and methods (Appendices 1)
Supplementary text (Appendices 2 to 5)
Figures S1 to S10
Tables S1 to S8

### Materials and Methods

### Appendix 1
### Damage function estimation and analysis

#### 1 Data source

##### 1.1 Damage function datasets

###### 1.1.1 Inclusion criteria

To estimate the damage function of individual episodes of selection $\tilde{f}(h)$, we used six criteria in our systematic search for appropriate datasets:

1. *Artificial experimental damage*

   We examined only studies that experimentally damaged the plants through artificial means (i.e., not with the use of herbivores, but scissors or clippers) because we can unambiguously attribute causation to the level of damage inflicted. Damage done with real herbivores suffer from bias because herbivores can adjust their feeding rate, which may be confounded with plant fitness, making our estimate of the damage function also biased. Although we acknowledge that the same level of physical damage may elicit different responses when the damage is made by real herbivores (*54*), damage made through artificial means is the only unbiased method to estimate plant tolerance to herbivore damage (*31*). Because we sought to estimate the specific shape of the damage functions, greater accuracy is more desirable.

2. *At least four levels of damage*

   We included only studies that manipulated damage at least four unique levels. This consideration is mainly in service of the identifiability of damage functions. With an intercept, our damage function has three degrees of freedom. Therefore, a minimum of four unique damage levels allows us to estimate the nonlinearity without parameter saturation.

3. *Independent means*

   Because we sought to estimate the nonlinearity in the damage functions, we excluded studies that reported group means based on an assumed functional form (typically linear or quadratic). Studies that reported raw mean plant responses or modeled mean plant responses from an ANOVA (therefore the means were estimated independently for each treatment level) were included.

4. *Damage levels can be expressed in % damage on the individual*

   Because we sought to characterize plant responses along $h$, which is in units of proportion damage, we excluded studies in which we could not express the damage levels as a percentage. Studies that cut plants to a specific height were included when the uncut plant height is known. Otherwise, it is difficult to translate the damage levels that can be related to theory.

5. *Measured fitness proxy*

   Because we sought to quantify how plant fitness responded to herbivore damage, we included only plant responses that can be interpreted in terms of fitness benefits or reduction (e.g., survival, germination, seed production, biomass production, seed mass, relative growth rate, flower production). We excluded chemical traits (e.g., C:N ratio, starch content, defensive chemistry concentration), ambiguous traits (e.g., specific leaf area, herbivory, root:shoot ratio), and phenology (e.g., day of flowering).

**Table S1: Paper filtering steps.** Numbers in parentheses are the number of papers. An addition sign '+' denotes when new papers were added to the filtering steps.

| Step (starting papers) | Exclusion (excluded papers) |
|---|---|
| Web of Science search (n = + 54,061) | Irrelevant document types (n = 539) |
| Filter by journal (n = 53,522) | Irrelevant journals (n = 3,017) |
| Filter by title & abstract (n = 50,505) | No keywords (n = 43,752) |
| Filter by paper metadata (n = 6,753) | Duplicated (n = 415) |
| Filter by abstract (n = 6,338) | No artificial damage (n = 4,886) |
| Filter by access (n = 1,452) | No access (n = 24) |
| Filter by quick skim (n = 1,428) | No $\geq$ 4 levels of damage (n = 1,169) |
| Add Google Scholar search (n = + 27) | |
| Filter by closer read (n = 286) | Cannot recover data (n = 6) |
| | Duplicated (n = 2) |
| | No $\geq$ 4 levels of damage (n = 70) |
| | No fitness proxy (n = 14) |
| | No independent means (n = 3) |
| | No percent damage (n = 57) |
| | No uncertainty (n = 17) |
| | Not experimental (n = 17) |
| Final inclusion (n = 100) | |

**Table S2: List of keywords used in the search.** A paper was included if it contains at least one of the keyword from each category. Variants of the same word were omitted for brevity.

| Category | Key words |
|---|---|
| Plant | plant, crop, tree, leaf, root, stem, seed, seedling, sapling, flower, fruit |
| Herbivore | herbivore, herbivory, grazing, grazer, browser, insect, pest, defoliator, hail |
| Response | fitness, yield, growth, seed, height, survival, reproduction, biomass, mass, weight |
| Treatment | damage, wound, clip, defoliate, removal, injury |
| Experiment | experiment, experimental, simulate, manipulate, artificial |

6. *Has uncertainty in means that can be recovered*

   Our data analysis pipeline requires uncertainty in the means for certain distributions. Without a measure of uncertainty, we were not able to parameterize distributions for parametric bootstrapping. For studies which did not report uncertainties, we tried to compute approximate uncertainties by using available information when possible (e.g., F-ratio, $r^2$, CV, P-values from F-test). We recovered these uncertainties from ANOVA type analysis by computing the residual standard deviation and used it as a measure of within-damage level uncertainty.

###### 1.1.2 Paper filtering

We searched for papers that matched our inclusion criteria on Web of Science and Google Scholar on 2024 January 31st. The filtering steps and sample sizes are shown in Table S1. We used the keywords shown in Table S2 in the initial search and in subsequent filtering steps programmatically. This is because we found that the search-engines did not abide by our filtering rules and the Web of Science database in particular, was missing a lot of abstracts. Missing abstracts were retrieved using the Crossref API (package RCROSSREF *55*).

##### 1.2 Species traits and study metadata

To examine whether plant characteristics explained the variation in damage function parameters, we collected experimental metadata from each publication and scored the relevant plant traits. For damaged organ, we classified it into five categories (root, stem/shoot, flower/seed/fruit, leaf, or unknown).

When leaves were damaged along with other organs, we scored the damaged organ based on the other organ. For damaged stage, we classified it into six categories (seed, seedling, vegetative, flowering, fruiting, or other). The 'other' category encompass experiments that damaged the plant at multiple different stages and studies that did not report the stage of damage. We also categorized tuber development as the 'other' category because the plant can be at multiple stages during tuber development. For domestication, we classified it into two categories (domesticated or wild). For growth form, we classified it into four categories (graminoid, climber, woody, or herbaceous). For life history, we classified it into three categories (growth, reproduction, or survival). For perenniality, we classified it into two categories (perennial or annual).

##### 1.3 Environmental predictors

To examine whether site characteristics explained the variation in damage function parameters, we retrieved existing environmental data using reported coordinates of where the artificial damage experiments were conducted (Table S3). Soil fertility is a qualitative ordered categorical variable which we analyzed as a numeric variable. The categories were divided into four groups corresponding to fertility that roughly allows for $< 40\%$, $40\% - 60\%$, $60\% - 80\%$, and $80\% - 100\%$ of the growth potential of plants. To obtain a site-level index of herbivory risk, we predicted mean plant-level herbivore damage from a Gradient Boosting Machine (GBM) model (package H2O *56*). GBM is an ensemble learning method that is effective at solving inverse-problems characterized by nonlinear relationships through sequentially fitting decision trees. We trained the model on two large global datasets of mean site-plant-level herbivore damage, 19 Bioclimatic variables, latitude, and elevation (Table S3). We used a 70/15/15 split, with 8,094 observations in the training dataset, 1,722 observations in the validation dataset, and 1,690 observations in the testing dataset. We did not include observations that were pooled across species. Observations of the same species at the same site were averaged. Hyperparameters were tuned based on 10-fold cross-validation and an early-stopping rule. The final model achieved a predictive performance of $r^2 = 0.37$ (mean absolute error = 0.088 proportion damage) on the testing dataset, indicating a reasonable ability to predict site-level herbivory risk.

**Table S3: Sources of data used in meta-regression.** NPP stands for annual Net Primary Productivity.

| Variable | Resolution | Use | Data source |
|---|---|---|---|
| BIO1-BIO19 (1970-2000) | 2.5' | Herbivory prediction | (*57*) |
| Elevation | 2.5' | Herbivory prediction | (*57*) |
| Mean herbivory | species-site | Herbivory prediction | (*29*)<br>(*58*) |
| Herbivory risk | 2.5' | Meta-regression | Predicted |
| Mean temperature (1970-2000) | 2.5' | Meta-regression | (*57*) |
| Mean precipitation (1970-2000) | 2.5' | Meta-regression | (*57*) |
| Mean NPP (2001-2010) | 15' | Meta-regression | (*59*) |
| Soil fertility | 0.5' | Meta-regression | (*60*) |

#### 2 Damage function estimation

##### 2.1 Episodes of selection

Our inability to have estimates of lifetime survival and fecundity, but only snapshots of their proxies, poses a fundamental challenge to estimate individual fitness and thus $f(h)$. To remedy this problem, we considered all fitness proxies as 'episodes of selection' (*sensu 33*, *61*). According to this assumption, the absolute fitness $W$ can be seen as an amalgamation of the fitness proxies in a series of $n$ multiplicative individual episodes of selection $\tilde{W}_i$,

$$W \equiv \prod_i^n \tilde{W}_i, \qquad n \geq 1, \tag{S1}$$

where we use the accent ˜ to denote variables in individual episodes of selection. This assumption implies that when we convert the fitness proxy measured in discrete time to continuous time by taking the logarithm, we can compare and interpret various fitness proxies by ignoring the intercept in our statistical models and the general baseline vigor of the plants considered (*62*). The additive decomposition also implies that by examining the average damage function of multiple episodes of selection $\langle \tilde{f}(h) \rangle = n^{-1} \sum_i^n \tilde{f}_i(h)$, we can approximate the shape of the damage function $f(h) \simeq n \langle \tilde{f}(h) \rangle$ and know that $|f(h)|$ must be at least as great as $|\langle \tilde{f}(h) \rangle|$, even if we do not know the exact magnitude. For conclusions that strictly rely on interpreting the average damage function parameters $\langle \tilde{\alpha} \rangle$ and $\langle \tilde{\beta} \rangle$, we implicitly assumed all else being equal.

#### 2.2 Function estimation

We fitted the parametric damage function $\tilde{f}(h; \tilde{\alpha}, \tilde{\beta}) = \tilde{\beta} h^{1/\tilde{\alpha}}$ to each dataset (Figure 1A). Let $\tilde{Y}$ be a fitness proxy of an episode of selection that is positively related to the overall plant fitness. We fitted the following model using a combination of optimization routines (package CALIBRAR *63*),

$$\ln\left(\tilde{Y}|h\right) \sim \mathcal{N}\left(\ln\left(\langle \tilde{Y}|h \rangle\right), \tilde{\epsilon}^2\right), \tag{S2a}$$

$$\ln(\langle \tilde{Y}|h \rangle) = \tilde{C} - \tilde{f}\left(h; \tilde{\alpha}, \tilde{\beta}\right). \tag{S2b}$$

Here, $\mathcal{N}$ denotes the normal distribution, $\tilde{\epsilon}^2$ denotes the error variance, and $\tilde{C}$ denotes some arbitrary constant for $\ln \tilde{Y}(h = 0)$, the average fitness proxy value when the plant receives no herbivory. For numerical stability when fitting (S2), we incorporated prior information to regularize the parameters of the fitted model. As such, the parameters were estimated via maximum *a posteriori* (MAP),

$$\underset{[\tilde{\alpha}, \tilde{\beta}, \tilde{C}, \tilde{\epsilon}^2]}{\arg\max}\ \mathcal{L}(\tilde{Y}|h; \tilde{\alpha}, \tilde{\beta}, \tilde{C}, \tilde{\epsilon}^2) \rho(\tilde{\alpha}, \tilde{\beta}, \tilde{C}, \tilde{\epsilon}^2), \tag{S3}$$

where $\mathcal{L}(\tilde{Y}|h; \cdot)$ is the likelihood function and $\rho(\cdot)$ is the prior distribution with hyper-parameters,

$$\ln(\tilde{\alpha}) \sim \mathcal{N}(0, \ln(3)), \tag{S4a}$$

$$\tilde{\beta} \sim \mathcal{N}(0, 3), \tag{S4b}$$

$$\tilde{C} \sim \mathcal{N}(0, 100), \tag{S4c}$$

$$\ln(\tilde{\epsilon}) \sim \mathcal{N}(0, \ln(3)). \tag{S4d}$$

### 3 Analysis pipeline

#### 3.1 Data preparation

We collected each dataset verbatim from studies and processed them with the following pipeline:

1. We performed data entry error checks, including implausible, impossible, and missing data values. Datasets without complete observations, including those missing measures of uncertainty were removed.

2. We matched each dataset to a pre-defined set of scores based on the recorded response variable name (e.g., seed production, plant survival, 1000 grain weight) and response unit (e.g., cm, count, log response ratio). The scores include the direction of the variable relative to plant fitness (positive or negative), the type of vital rate (growth, reproduction, survival), the conditional distribution of the variable (log-normal, normal, Poisson, binomial, gamma, beta), and necessary back-transformations (e.g., undo square-root transformations on counts, convert percent to proportions, convert percent change to log-response ratio). Given that the log-normal distribution is not a member of the exponential dispersion family, we log-transformed all log-normal datasets and set their conditional distribution as normal. For count data which we have estimates of uncertainty, we used the gamma distribution, as it is a continuous approximation of the negative binomial distribution which allows the variance to be modeled independently of the mean. When uncertainty was not available for count data, we assumed that the data is Poisson distributed and therefore no variance estimate was needed.

3. We transformed the observed means and uncertainty measures of each damage level according to the corresponding back-transformation. The uncertainty was converted to standard errors. We used the delta method to approximate the variance-covariance matrix of the transformed random variables.

4. We performed 200 parametric bootstraps of each damage level using the mean, standard error, and sample size to include data uncertainty in our model. For datasets which we can access the raw data (as opposed to the summarized group means), we performed the same bootstrap procedure, but setting the standard error to zero. For datasets with a response variable that is beta or binomially distributed, we simulated the bootstrap datasets directly from the conjugate posterior distribution, the beta distribution with shape parameters $\alpha$ and $\beta$. We used a Bayes-Laplace prior $\mathcal{B}(\alpha = 1, \beta = 1)$, which places uniform density on all proportions (*64*). Therefore, for the posterior distribution of the mean, we write,

$$\mathcal{B}(\alpha = 1 + \text{N observed successes}, \beta = 1 + \text{N observed failures}). \tag{S5}$$

   Incorporating prior information in the bootstrap datasets avoided a lot of problems introduced by finite sample sizes. For instance, with only three observations, a sample is very likely to contain all 0's or 1's, which has a sample standard error of 0. Of course, we do not actually believe with infinite certainty that the true proportion is 0 or 1 based on three observations. Likewise, with limited observations of a Poisson distribution, we seldom actually believe that the true rate is 0. Therefore, for datasets with a response variable that is Poisson distributed, we simulated the bootstrap datasets directly from the conjugate posterior distribution, the gamma distribution with shape parameter $\alpha$ and rate parameter $\beta$ respectively. We used a uniform prior $\mathcal{G}(\alpha = 1, \beta = 0)$ (*64*), so that the posterior distribution of the mean reads,

$$\mathcal{G}(\alpha = 1 + \text{sum of observed counts}, \beta = \text{N observations}). \tag{S6}$$

   These conjugate distributions have the important property of regularizing the posterior mean when the sample size is small, but allowing the posterior mean to converge to the mean of the sample when the sample size is large.

5. We transformed each bootstrap datasets to be positively related to plant fitness.

6. We fitted the applicable model (S2, S41, S42, S43) to each bootstrap dataset.

7. We extracted the fitted model coefficients for subsequent analyses if applicable.

8. We simulated 200 damage functions from each fitted model by drawing fitted coefficients from a multivariate normal distribution $\mathbf{\Theta} \sim \mathcal{N}(\hat{\mathbf{\Theta}}, \hat{\Sigma}_{\mathbf{\Theta}})$. After that, we used these damage functions to compute the selection gradient and fitness differential. We estimated the Malthusian fitness differential in an episode of selection $\Delta\tilde{w}(h)$, where $w \propto \ln R_0 = \ln W$, and the fitness gradient in an episode of selection $\nabla\tilde{w}(h)$ evaluated at different locations along a gradient of herbivory intensity $h \in \{0.05, 0.15, 0.25, 0.5, 0.75, 0.95\}$. These specific locations were chosen somewhat arbitrarily but are representative of different regions of the damage functions. The first two locations in particular correspond roughly to the low and high end of average herbivore damage observed in field studies (*29*, *65*, *66*). The computation of the fitness differential $\Delta\tilde{w}(h)$ relative to the control (with no damage) was fairly straightforward once the damage function was estimated and noting that $\tilde{f}(0) = 0$,

$$\Delta\tilde{w}(h) = -\tilde{f}(h) + \tilde{f}(0) = -\tilde{f}(h) \tag{S7}$$

   Likewise, for the fitness gradient $\nabla\tilde{w}(h)$, we used the forward finite difference method with the step size $d = 0.01$,

$$\nabla\tilde{w}(h) \approx -\frac{\tilde{f}(h+d) - \tilde{f}(h)}{d}. \tag{S8}$$

9. For a collection of estimates of interest $X_{i,j}$ generated from the $i$th draw of a model fitted to the $j$th bootstrap dataset, we pooled the estimates as a mean estimate $\hat{x}$ and standard error $\hat{s}$ given

by,

$$\hat{x} = \frac{1}{m}\sum_{j=1}^{m}\left(\frac{1}{n}\sum_{i=1}^{n} X_{i,j}\right), \tag{S9a}$$

$$\hat{s} = \sqrt{\underbrace{\langle \sigma_j^2(\mathbf{X})\rangle}_{\text{Model uncertainty}} + \underbrace{\sigma^2(\langle \mathbf{X}\rangle_j)}_{\text{Data uncertainty}}}, \tag{S9b}$$

where $\mathbf{X}$ is an $n \times m$ matrix. We found that data uncertainty and model uncertainty generally contribute in equal proportion to total pooled uncertainty.

##### 3.2 Damage function parameter classification

The high heterogeneity in individual damage functions we observed raises the question of how often different damage function responses are exhibited at the individual function level. To address this question, we needed to classify damage function parameters into different qualitative categories (e.g., sensitive, neutral, insensitive) and ask what proportion of damage functions fall under each category. Accordingly, we used adaptive shrinkage, an empirical Bayes approach from (package ASHR *67*, *68*). The method enables information sharing across individual damage function estimates and provides a straightforward way to summarize posterior probabilities globally, substantially reducing uncertainty in the overall proportion of classifications while maintaining control of the false discovery rate. Specifically, we assume that a true parameter value $x_j$ is drawn from a generic unimodal prior distribution which is centered at 0 and may be asymmetrical. It consists of a mixture between a point mass on the null hypothesis $x = 0$ and a distribution of the alternate hypothesis approximated as a mixture of half uniform distributions. The posterior distribution $P(x; \hat{x}, \hat{s})$ under these assumptions can be used to compute the posterior probability that a given true parameter value fall above or below some threshold, providing a means to estimate the global share of different classifications. For example, to compute the estimated proportion of damage functions classified as sensitive, neutral, or insensitive (letting $\hat{x} \equiv \ln \tilde{\alpha}$), we used $\langle P(x > 0; \hat{x}, \hat{s})\rangle$, $\langle P(x = 0; \hat{x}, \hat{s})\rangle$, and $\langle P(x < 0; \hat{x}, \hat{s})\rangle$ respectively.

##### 3.3 Estimation of group means

###### 3.3.1 Damage function parameters

To estimate how damage function parameters vary across environmental gradients and plant functional groups, we fitted multilevel random-effects models with the following structure (package METAFOR *69*),

$$\begin{aligned}
\mathbf{y} &\sim \mathcal{N}(\bar{\mathbf{y}}, \boldsymbol{\varepsilon}^{\mathbf{2}}\mathbf{I}_{\text{eff}}),\\
\bar{\mathbf{y}} &= \mathbf{X}\mathbf{b}\\
&\quad + \mathbf{Z}_{\text{sp}}\mathbf{a}_{\text{sp}} + \mathbf{Z}_{\text{sp}}\mathbf{a}_{\text{phy}}\\
&\quad + \mathbf{Z}_{\text{pub}}\mathbf{a}_{\text{pub}} + \mathbf{Z}_{\text{exp}}\mathbf{a}_{\text{exp}} + \mathbf{Z}_{\text{eff}}\mathbf{a}_{\text{eff}},\\
\mathbf{a}_{\text{sp}} &\sim \mathcal{N}(\mathbf{0}, \sigma^2_{\text{sp}}\mathbf{I}_{\text{sp}}),\\
\mathbf{a}_{\text{phy}} &\sim \mathcal{N}(\mathbf{0}, \sigma^2_{\text{phy}}\boldsymbol{\Sigma}_{\text{phy}}),\\
\mathbf{a}_{\text{pub}} &\sim \mathcal{N}(\mathbf{0}, \sigma^2_{\text{pub}}\mathbf{I}_{\text{pub}}),\\
\mathbf{a}_{\text{exp}} &\sim \mathcal{N}(\mathbf{0}, \sigma^2_{\text{exp}}\mathbf{I}_{\text{exp}}),\\
\mathbf{a}_{\text{eff}} &\sim \mathcal{N}(\mathbf{0}, \sigma^2_{\text{eff}}\mathbf{I}_{\text{eff}}).
\end{aligned} \tag{S10}$$

Here, $\mathbf{y}$ denotes a vector of point estimate of the response variable, with total measurement uncertainty as $\boldsymbol{\varepsilon}^{\mathbf{2}}$ from (S9). We used $\mathbf{X}$ and $\mathbf{Z}$ to denote the design matrices, $\mathbf{I}$ to denote the identity matrix, and $\mathbf{b}$ and $\mathbf{a}$ to denote fixed and random-effects, respectively. We accounted for non-independence at the species level by estimating the variance (heterogeneity) parameters $\sigma^2_{\text{sp}}$ and $\sigma^2_{\text{phy}}$ for species identity and species phylogenetic relatedness respectively. We derived the species variance-covariance matrix $\boldsymbol{\Sigma}_{\text{phy}}$ using (package PHYTOOLS *70*) and a phylogenetic tree (Figure S5) pruned from the most recent angiosperm megaphylogeny (package RTREES *71*–*73*). We also accounted for publication and experimental identity level non-independence by estimating separate variance parameters $\sigma^2_{\text{pub}}$ and

$\sigma^2_{\text{exp}}$ respectively for each level. All sources of variation within an experiment was estimated as an individual effect level heterogeneity $\sigma^2_{\text{eff}}$. The details of the fixed effects depended on the specific model, but all models included an intercept term (Table S4). For the analysis of the parameter $\tilde{\alpha}$, we estimated $\tilde{\alpha}|\tilde{\beta} > 0$ and $\tilde{\alpha}|\tilde{\beta} < 0$ separately via no-pooling by including a binary variable for whether $\tilde{\beta}$ is positive (i.e., damaging). We analyzed the parameter separately because meaningful biological interpretation of $\tilde{\alpha}$ is dependent on whether tissue damage is damaging or benefiting. To get a case specific measure of predictive performance (i.e., AICc), we also fitted $\tilde{\alpha}|\tilde{\beta} > 0$ and $\tilde{\alpha}|\tilde{\beta} < 0$ separately in two additional sets of models. We did not consider the effect sizes estimated from these models as they would be inferior to that of the full model that has access to more information. Finally, for the analysis of site characteristics, we included only studies which were conducted in the field (i.e., excluded laboratory and greenhouse studies).

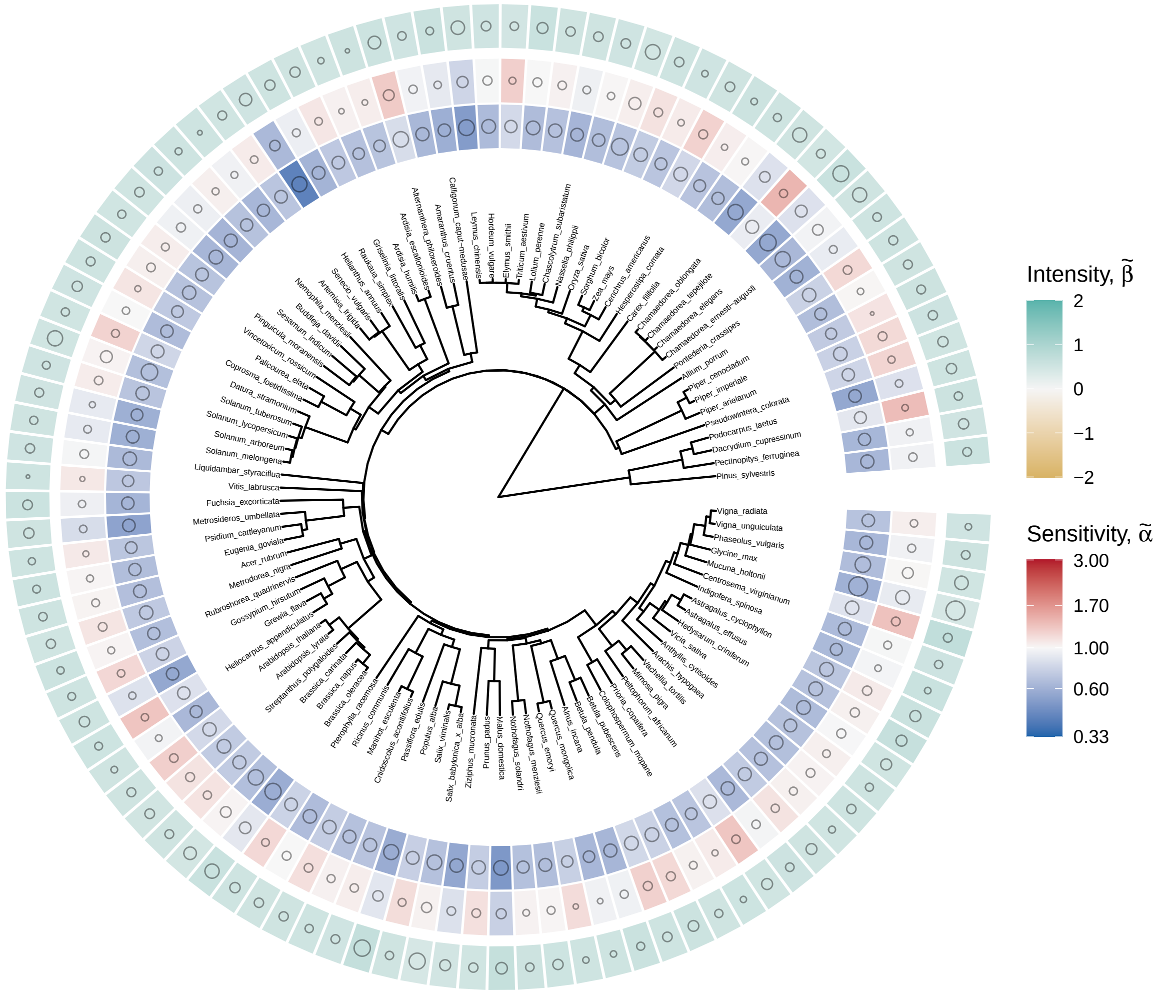

**Figure S5: The estimated damage function parameters are distributed randomly with respect to the phylogeny of species included in the analysis.** The concentric circles from innermost to outermost show $\tilde{\alpha}|\tilde{\beta} > 0$, $\tilde{\alpha}|\tilde{\beta} < 0$, and $\tilde{\beta}$ respectively. The size of each open circle in each tile indicates the precision of the estimate (1/SE). Estimates were predicted from a fitted random-effect model. There is no evidence of phylogenetic signal for any of the three variables shown here (Pagel's $\lambda \simeq 0$, $P \simeq 1$) (package PHYTOOLS *70*).

##### 3.3.2 Fitness differential and gradient

To estimate the average fitness differential and gradient, we used a similar model structure as (S10) but with some small modifications. First, we included a random intercept $\mathbf{a}_{\text{dataset}}$ at the individual damage function level because we derived multiple effect sizes from the same dataset. Second, we dropped the

**Table S4: Specification of fixed effects across all multi-level random-effects models.** We used 'predictor' as a place holder for different predictors considered in different model sets (plant characteristics: {damaged organ, damaged stage, domesticated, growth form, life history, perenniality}; site characteristics and predictive performance: {absolute latitude, net primary productivity, mean annual temperature, mean annual precipitation, herbivory risk, soil fertility}). Each predictor is fitted in a different model. We used 'ns()' to denote natural cubic splines with three degrees of freedom (package SPLINES 74). Random effects are described in the text.

| Model sets | Response variable | Fixed effects |
|---|---|---|
| Plant characteristics | $\tilde{\alpha}$ | Intercept + predictor × damaging |
| Plant characteristics | $\tilde{\beta}$ | Intercept + predictor |
| Site characteristics | $\tilde{\alpha}$ | Intercept + predictor × damaging × life_history |
| Site characteristics / predictive performance | $\tilde{\beta}$ | Intercept + predictor × life_history |
| Predictive performance | $\tilde{\alpha}\|\tilde{\beta} > 0$ | Intercept + predictor × life_history |
| Predictive performance | $\tilde{\alpha}\|\tilde{\beta} < 0$ | Intercept + predictor × life_history |
| – | Fitness differential | ns($h$) |
| – | Fitness gradient | Intercept + ns($h$) |

phylogenetic term to take advantage of METAFOR's sparse matrix routine, which dramatically reduced computational time. This move was justified because phylogenetic signals is generally very weak in our data (Figure S5). The details of the fixed effects again depended on the specific model (Table S4). We did not estimate an intercept for fitness differential because we know that it should be 0 when there is no damage (i.e., $\langle \tilde{f}(0) - \tilde{f}(0) \rangle = 0$). Because we expect that the model relationship between the response variable and proportion damage is nonlinear, we modeled the relationship with cubic splines. Our full model thus reads,

$$
\begin{aligned}
\mathbf{y} &\sim \mathcal{N}(\bar{\mathbf{y}}, \boldsymbol{\varepsilon}^2 \mathbf{I}_{\text{eff}}), \\
\bar{\mathbf{y}} &= \mathbf{X}\mathbf{b} \\
&\quad + \mathbf{Z}_{\text{sp}}\mathbf{a}_{\text{sp}} \\
&\quad + \mathbf{Z}_{\text{pub}}\mathbf{a}_{\text{pub}} + \mathbf{Z}_{\text{exp}}\mathbf{a}_{\text{exp}} + \mathbf{Z}_{\text{dataset},}\mathbf{a}_{\text{dataset}} + \mathbf{Z}_{\text{eff},}\mathbf{a}_{\text{eff}}, \\
\mathbf{a}_{\text{sp}} &\sim \mathcal{N}(\mathbf{0}, \sigma^2_{\text{sp}} \mathbf{I}_{\text{sp}}), \\
\mathbf{a}_{\text{pub}} &\sim \mathcal{N}(\mathbf{0}, \sigma^2_{\text{pub}} \mathbf{I}_{\text{pub}}), \\
\mathbf{a}_{\text{exp}} &\sim \mathcal{N}(\mathbf{0}, \sigma^2_{\text{exp}} \mathbf{I}_{\text{exp}}), \\
\mathbf{a}_{\text{dataset}} &\sim \mathcal{N}(\mathbf{0}, \sigma^2_{\text{dataset}} \mathbf{I}_{\text{dataset}}), \\
\mathbf{a}_{\text{eff}} &\sim \mathcal{N}(\mathbf{0}, \sigma^2_{\text{eff}} \mathbf{I}_{\text{eff}}).
\end{aligned}
\tag{S11}
$$

All analyses are conducted in R version 4.4.2 (74).

### Supplementary Text

### Appendix 2
### Population dynamics

#### 4 Introduction

Here, we examine how the shape of the herbivory damage function affects the population dynamics of herbivores and plants. We consider a relatively simple two-species plant-herbivore model based on the classic work of Crawley (*21*) (hereafter 'CR'), and Anderson and May (*75*, *76*) (hereafter 'AM'). Our goal is to derive a new set of stability criteria and expressions for equilibria based on the family of damage functions (a generalization of the special cases CR and AM analyzed) we explored in the main text. We begin by briefly summarizing the rationale behind the model construction before considering special cases where we explore nonlinearity in damage to plant reproduction and plant survival. In particular, we address the question of how nonlinearity in the herbivory damage function affects the value and stability of the non-trivial equilibrium, and whether the herbivores can regulate (exert top-down control on) the plant population.

##### 4.1 General model components

The CR and AM model of plant-herbivore dynamics considers a population of herbivores $N$ and plants $V$ where the population-level outcomes of plant-herbivore interactions are modeled as the expectation of local interactions on individual plants. The local interactions are dependent on the local number of resident herbivores $n$, which is drawn from a probability mass function $P(n)$, but are independent from interactions occurring on other plant individuals. The model assumes that herbivore death, plant death, and plant reproduction can occur at some intrinsic rate. Herbivores are born as a result of plant consumption but which may reduce the fecundity and survival of host plants. Additionally, herbivores may die when their host plant dies.

We modified this model to accommodate the damage function $f(h)$. Writing the proportional effect of damage $h$ on individual plant reproduction $b$ or mortality $\mu_V$ as $F(h)$ (that is, $bF_b(h) = f_b(h)$ and $\mu_V F(h) = f_\mu(h)$), we additionally introduce the density dependent herbivore damage function $h(n)$ to relate the local herbivore density $n$ to the average population-level relative effect on vital rate,

$$\sum_{n=0}^{\infty} F(h(n))P(n). \tag{S12}$$

Then, defining the herbivore transmission success function as $\rho(V)$ and the density dependent herbivore

**Table S5: Principal variables considered in the model.**

| Class | Symbol | Definition | Unit |
|---|---|---|---|
| state variable | $V$ | Plant density | number of plants |
| state variable | $N$ | Herbivore density | number of herbivores |
| state variable | $\lambda$ | Average herbivore load | average herbivore per plant |
| state variable | $h$ | herbivore damage | proportion |
| functional response | $F(h;\alpha,\beta)$ | damage function (total fitness) | time$^{-1}$ |
| parameter | $\alpha$ | damage sensitivity (total fitness) | unitless |
| parameter | $\beta$ | damage intensity (total fitness) | ratio of $\mu_V$ or $b$ per proportion of damage* |
| parameter | $b$ | per-capita plant birth rate | time$^{-1}$ |
| parameter | $\mu_V$ | per-capita plant death rate | time$^{-1}$ |
| parameter | $\mu_N$ | per-capita herbivore death rate | time$^{-1}$ |
| parameter | $\delta$ | per-capita plant consumption | proportion of damage herbivore$^{-1}$ |
| parameter | $c$ | conversion efficiency | herbivore time$^{-1}$plant$^{-1}$ |

*Note that $\beta$ and $\tilde{\beta}$ presented in the main text and elsewhere besides Appendix 2 subsume $b$ and $\mu_V$ and have the unit time$^{-1}$ per proportion of damage.

mortality rate as $m(n)$, the plant-herbivore dynamics can be generically written as (Table S5),

$$\frac{dV}{dt} = \underbrace{Vb}_{\substack{\text{Intrinsic}\\ \text{births}}} - \underbrace{Vb\left(\sum_{n=0}^{\infty} F_b\left(h(n)\right)P(n|\lambda)\right)}_{\text{Herbivore induced reduction in fecundity}} - \underbrace{V\mu_V}_{\substack{\text{Intrinsic}\\ \text{plant deaths}}}$$

$$- \underbrace{V\mu_V\left(\sum_{n=0}^{\infty} F_\mu\left(h(n)\right)P(n|\lambda)\right)}_{\text{Herbivore induced reduction in survival}}, \tag{S13a}$$

$$\frac{dN}{dt} = \underbrace{Vc\left(\sum_{n=0}^{\infty} h(n)P(n|\lambda)\right)\rho(V)}_{\text{Herbivore births from feeding}} - \underbrace{N\mu_V}_{\substack{\text{Intrinsic}\\ \text{plant deaths}}}$$

$$- \underbrace{V\mu_V\left(\sum_{n=0}^{\infty} F_\mu\left(h(n)\right)P(n|\lambda)n\right)}_{\substack{\text{Herbivore death due to herbivore}\\ \text{induced plant deaths}}} - \underbrace{V\left(\sum_{n=0}^{\infty} m(n)P(n|\lambda)\right)}_{\text{Herbivore intrinsic deaths}}, \tag{S13b}$$

where $\lambda$ is the average herbivore load,

$$\lambda \equiv \frac{N}{V} = \langle n \rangle. \tag{S14}$$

We used $\langle \cdot \rangle$ to denote the mean and will use this notation throughout. Note that following AM and CR, we assume plants grow exponentially in the absence of herbivores. This simplifying assumption lets us isolate the effects of nonlinear damage functions on dynamics without the complicating factor of density-dependent plant growth.

#### 4.2 Further model assumptions

Now suppose for simplicity that:

1. The herbivores are Poisson distributed on individual plants, so that,

$$P(n|\lambda) \equiv \frac{\lambda^n e^{-\lambda}}{n!}. \tag{S15}$$

   This is one of the cases examined by both CR and AM and has the advantage of being simple. (*30*) found that this assumption approximates the distribution of herbivory among plants quite well.

2. Herbivore damage increases linearly with local herbivore density,

$$h(n;\delta) \equiv \delta n, \tag{S16}$$

so that each herbivore individual eats the same amount $\delta$. Although technically this assumption means that the herbivores can eat more than the available plant biomass when the local herbivore density is very high, as we shall discuss later, we impose a feasibility condition to avoid this problem. Such simplification is necessary for us to focus on the effect of nonlinearity introduced by the damage function $f(h)$, as opposed to nonlinearity introduced by herbivory.

3. The herbivore transmission success increases linearly with plant density,

$$\rho(V) \equiv V. \tag{S17}$$

This rate of increase for herbivores follows CR, a departure from the type II functional response in AM.

4. The per capita rate of death of herbivores increases quadratically with density,

$$m(n;\mu_N) \equiv \mu_N n^2. \tag{S18}$$

The quadratic term introduces linear density dependence for herbivores per one of the special cases examined by AM. We included density dependence to prevent herbivores from often growing unregulated, a side effect of CR's modification to the transmission success function.

5. The scaled damage function (on either plant reproduction or survival) follows the form (Figure 1A),

$$F(h;\alpha,\beta) \equiv \beta h^{1/\alpha}. \tag{S19}$$

Because we scaled the damage function proportional to the intrinsic growth rate and mortality rate of the plant such that $\beta_b \in [0,\infty]$ and $\beta_\mu \in [0,\infty]$, the intensity parameter $\beta$ and can be interpreted as the proportional change in reproduction or mortality rate. We could in theory extend the support of $\beta$ to include negative numbers, as in the main text, whereby plant fitness is benefited by herbivore damage. However, given that the plant dynamics is not regulated by any other factor by design, without damaging herbivores (where $\beta \leq 0$), the plant population simply exhibits unbounded exponential growth, at least until it is limited by other factors. $\alpha \in (0,\infty)$ is a shape parameter that controls the convexity of the damage function. When $\alpha < 1$, the damage function is convex with respect to plant fitness, so that additional damage reduces plant fitness more; when $\alpha > 1$ the function is concave with respect to plant fitness, so that additional damage reduces plant fitness less; when $\alpha = 1$ the function is linear, and additional damage reduces plant fitness by the same amount regardless of preexisting damage. The power damage function approximately captures the range of shapes that can be found in nature as well as the shape of more mechanistic models that consider photosynthesis (*15*) or smoothed transition between limiting resources (*36*). Similar range of damage function forms have been proposed qualitatively in (*16*) and (*77*), and quantitatively in (*78*) (although no model analysis was presented).

Substituting the above definitions, and noting that $V\sum_{n=0}^{\infty} nP(n|\lambda) = V\lambda = N$ and $\sum_{n=0}^{\infty} n^2 P(n|\lambda) = \lambda^2 + \lambda$, yields,

$$\frac{dV}{dt} = Vb\left(1 - \beta_b \delta^{1/\alpha_b} \sum_{n=0}^{\infty} n^{1/\alpha_b} P(n|\lambda)\right) - V\mu_V\left(1 + \beta_\mu \delta^{1/\alpha_\mu} \sum_{n=0}^{\infty} n^{1/\alpha_\mu} P(n|\lambda)\right), \tag{S20a}$$

$$\frac{dN}{dt} = N\left(c\delta V - \mu_V\right) - V\left(\mu_V\left(\beta_\mu \delta^{1/\alpha_\mu} \sum_{n=0}^{\infty} n^{1/\alpha_\mu + 1} P(n|\lambda)\right) + \mu_N(\lambda^2 + \lambda)\right). \tag{S20b}$$

The sums can be seen as raw moments of the Poisson distribution. For integer moments, we have exactly,

$$\langle n^{1/\alpha} \rangle = \sum_{i=0}^{1/\alpha} \lambda^i s(1/\alpha, i), \tag{S21}$$

where $s(1/\alpha, i)$ is the Stirling number of the second kind. For non-integer fractional moments, there is no closed-form expression, but we can use the approximation for large $\lambda$, where $\langle n^{1/\alpha} \rangle \simeq \lambda^{1/\alpha}$, allowing us to write,

$$\frac{dV}{dt} = V\left(b(1 - \beta_b(\delta\lambda)^{1/\alpha_b}) - \mu_V(1 + \beta_\mu(\delta\lambda)^{1/\alpha_\mu})\right), \tag{S22a}$$

$$\frac{dN}{dt} = N\left(c\delta V - \mu_V(1 + \beta_\mu(\delta\lambda)^{1/\alpha_\mu}) - \mu_N(\lambda + 1)\right). \tag{S22b}$$

In the following, we analyzed special cases of this general representation.

### 5 Model analysis

#### 5.1 Reduction in fecundity

We assessed how the shape of the damage function affects the equilibrium and stability of the plant-herbivore dynamics when herbivores only affect plant reproduction. When the damage of herbivores is restricted to host reproduction, we can set $\beta_\mu = 0$, yielding,

$$\frac{dV}{dt} = V\left(b\left(1 - \beta(\delta\lambda)^{1/\alpha}\right) - \mu_V\right), \tag{S23a}$$

$$\frac{dN}{dt} = N\left(c\delta V - \mu_V - \mu_N(\lambda + 1)\right). \tag{S23b}$$

The subscripts were removed for simplicity. Notice that if we just set the second term in (S22b) to zero, and either assume $\alpha_\mu = \alpha_b$ or $\beta_b = 0$ in (S22a), the model reduces to a special case of (S23). In other words, if we relax the assumption that the herbivores die when they kill their host plant, the dynamics resemble a reduction in fecundity regardless of whether herbivores affect plant fitness through fecundity or mortality.

##### 5.1.1 Equilibrium

Next, we find the non-trivial equilibrium. Setting (S23a) to 0 and $V \neq 0$, and solving for $\lambda^*$, we have the non-trivial equilibrium mean herbivore load of each plant,

$$\lambda^* = \frac{1}{\delta}\left(\frac{1 - \frac{\mu_V}{b}}{\beta}\right)^\alpha. \tag{S24}$$

Setting (S23b) to 0 and $N \neq 0$, then recalling (S14), lead to the non-trivial population equilibrium,

$$V^* = \frac{1}{c\delta}\left(\mu_V + \mu_N(\lambda^* + 1)\right), \tag{S25a}$$

$$N^* = \frac{\lambda^*}{c\delta}\left(\mu_V + \mu_N(\lambda^* + 1)\right). \tag{S25b}$$

Reassuringly, the positive relationship between herbivore load and plant density implied by (S25) is consistent with the Resource Concentration Hypothesis (*79*), which enjoys wide empirical support (*80*). For the above equilibrium to be feasible, the equilibrium mean herbivore load must be positive; this leads to the condition that the net intrinsic growth rate of the plant must be positive,

$$\lambda^* > 0 \iff b > \mu_V. \tag{S26}$$

Likewise, the amount of herbivore damage on a single plant, at least on average, cannot exceed unity based on our definition that $h \in [0, 1]$ is a proportion, leading to an upper bound for the plant intrinsic birth rate $b$,

$$\begin{aligned} h(\lambda^*) \leq 1 &\iff \delta\lambda^* \leq 1, \\ &\iff b \leq \frac{\mu_V}{1 - \beta}. \end{aligned} \tag{S27}$$

With the bound set by (S27), we can find that the quantity inside the parentheses of (S24) is strictly less than or equal to 1. This implies that $\lambda^*$, and, by extension, $V^*$ and $N^*$ are all decreasing functions of $\alpha$ and $\beta$ (Figure S6). In other words, the less tolerant the plants are to herbivore damage, the lower mean herbivore load, herbivore density, and plant density can be maintained. This result makes intuitive sense because greater damage tolerance at low-damage levels and lower fitness reduction at high-damage levels allow plants to achieve a higher population. With a greater population of plants to exploit and with more tolerant plant individuals which can sustain a higher herbivore load, the herbivore population also increases.

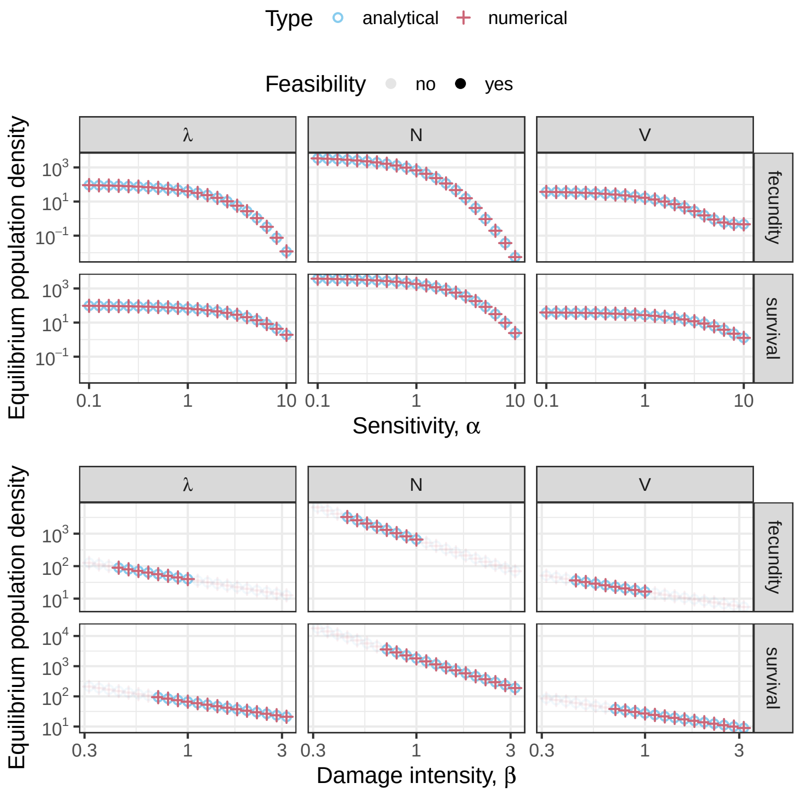

**Figure S6: The non-trivial equilibrium value for herbivore density $N$, herbivore load $\lambda$, and plant density $V$ when herbivore damage reduces fecundity or survival.** (A) slice through the damage function shape parameter space $\alpha$. (B) slice through the damage function intensity parameter space $\beta$. Analytical values (purple cross) come from (S24), (S25), (S32), and (S33). Numerical values (blue circles) come from numerically solving (S23) or (S31) (package deSolve *81*). Infeasible equilibria have a higher transparency. We used the values $\delta = 0.01, b = 0.1, \mu_V = 0.06, \mu_N = 0.4, c = 100$ for both (A) and (B). In (A), $\beta = 0.99$. In (B), $\alpha = 1$.

##### 5.1.2 Stability and regulation

Next, we found the stability criteria of (S25). The above model has the Jacobian,

$$\mathbf{J}(V,N) = \begin{pmatrix} (\frac{1}{\alpha}-1)\beta b\,(\delta\lambda)^{1/\alpha} - \mu_V + b & -\frac{\beta b}{\alpha}\,(\delta\lambda)^{1/\alpha}\,\lambda^{-1} \\ Nc\delta + \lambda^2\mu_N & c\delta V - \mu_N(2\lambda+1) - \mu_V \end{pmatrix}. \tag{S28}$$

Using the Routh–Hurwitz stability criterion, we find the stability conditions for the non-trivial equilibrium,

$$0 > \frac{b-\mu_V}{\alpha} - \lambda^*\mu_N, \tag{S29a}$$

$$0 < \frac{b-\mu_V}{\alpha}(\mu_V + \mu_N(\lambda^*+1)). \tag{S29b}$$

Recalling the feasibility condition (S26) allows us to find that $(b - \mu_V) > 0$ and therefore (S29b) is always satisfied. Leaving us with the final stability and feasibility conditions,

$$\mu_V < b \leq \frac{\mu_V}{1-\beta}, \tag{S30a}$$

$$\lambda^* \mu_N > \frac{b - \mu_V}{\alpha}. \tag{S30b}$$

We can see here that increasing $\beta$ always decreases the stability of the equilibrium, although it increases the feasibility of the equilibrium (Figure S7). This result is qualitatively consistent with the finding of AM. For $\alpha$, the result is more complicated because increasing values of $\alpha$ decreases both sides of the inequality (S30b). Intermediate values of $\alpha$ is the most stable, and though stability decreases with increasingly extreme sensitivity or insensitivity, so long that the net plant growth rate is not too large (a likely scenario in nature), increasing insensitivity (decreasing $\alpha$) allows the system to be stable (Figure S7).

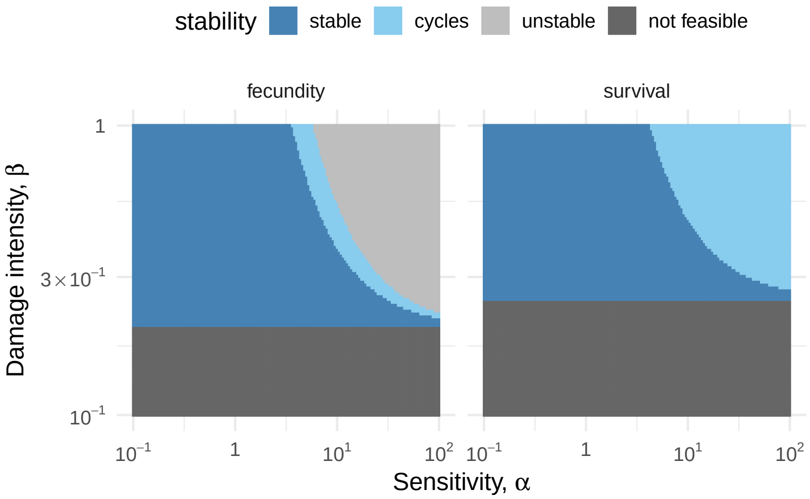

**Figure S7: Stability of the non-trivial equilibrium when herbivore damage reduces plant fecundity or survival.** Results based on Routh–Hurwitz stability criterion. We used the values $\delta = 0.01, b = 0.1, \mu_V = 0.08, \mu_N = 0.4, c = 100$. The system would cycle if $\det(\mathbf{J}(V^*, N^*)) > \frac{1}{4}\mathrm{Tr}(\mathbf{J}(V^*, N^*))^2$ and eventually converge to the equilibrium.

#### 5.2 Reduction in survival

Here, we examine when herbivory only reduces plant survival by setting $\beta_b = 0$ in (S22), which now reads,

$$\frac{dV}{dt} = V\left(b - \mu_V(1 + \beta(\delta\lambda)^{1/\alpha})\right), \tag{S31a}$$

$$\frac{dN}{dt} = N\left(c\delta V - \mu_V(1 + \beta(\delta\lambda)^{1/\alpha}) - \mu_N(\lambda + 1)\right). \tag{S31b}$$

Again, we dropped the subscripts here for simplicity.

##### 5.2.1 Equilibrium

Setting (S31a) and $V \neq 0$, then solving for $\lambda^*$, we have,

$$\lambda^* = \frac{1}{\delta}\left(\frac{\frac{b}{\mu_V} - 1}{\beta}\right)^{\alpha}. \tag{S32}$$

Setting (S31b) and $N \neq 0$, then using (S14), we have,

$$V^* = \frac{1}{c\delta}\left(b + \mu_N(\lambda^* + 1)\right), \tag{S33a}$$

$$N^* = \frac{\lambda^*}{c\delta}\left(b + \mu_N(\lambda^* + 1)\right). \tag{S33b}$$

For the equilibrium to be feasible, we have the same criterion as earlier,

$$\lambda^* > 0 \iff b > \mu_V. \tag{S34}$$

In addition, for herbivore damage to be feasible, we have another feasibility criterion,

$$\begin{aligned} h(\lambda^*) \leq 1 &\iff \delta\lambda^* \leq 1, \\ &\iff \frac{b}{\mu_V} \leq \beta + 1. \end{aligned} \tag{S35}$$

With the bound set by (S35), we can find that the quantity in the inner parentheses of (S32) is strictly less than or equal to 1. Therefore, we have similar to the case if herbivore damage reduces plant fecundity that $\alpha$ decreases the equilibrium herbivore load $\lambda^*$, and by extension the herbivore $N^*$ and plant population density $V^*$ (Figure S6). The same qualitative result is found by CR and AM who examined a narrower range of $\alpha \in \{1, \frac{1}{2}, \frac{1}{3}\}$. Finally, we can see that greater $\beta$ increases the feasibility of the equilibrium, but decreases the equilibrium mean parasite load, plant population density, and herbivore population density.

##### 5.2.2 Stability and regulation

We derived the stability criteria of (S33). The above model has the Jacobian,

$$\mathbf{J}(V,N) = \begin{pmatrix} (\frac{1}{\alpha} - 1)\beta\mu_V\,(\delta\lambda)^{1/\alpha} - \mu_V + b & -\frac{\beta\mu_V}{\alpha}\,(\delta\lambda)^{1/\alpha}\,\lambda^{-1} \\ Nc\delta + \lambda^2\mu_N + \frac{\beta\lambda\mu_V}{\alpha}(\delta\lambda)^{1/\alpha} & c\delta V - \mu_N(2\lambda+1) - \mu_V\left((\frac{1}{\alpha}+1)\beta(\delta\lambda)^{1/\alpha} + 1\right) \end{pmatrix}, \tag{S36}$$

and stability conditions at the non-trivial equilibrium,

$$0 > -\mu_N\lambda^*, \tag{S37a}$$

$$0 < \frac{b - \mu_V}{\alpha}(b + \mu_N\,(\lambda^* + 1)). \tag{S37b}$$

Recall from (S34) that $(b - \mu_V) > 0$, so long as $\mu_N > 0$ (when $\mu_N = 0$, the system is neutrally stable), both conditions can be satisfied; therefore, the simplified stability and feasibility criteria read,

$$\mu_V < b \leq \mu_V(\beta + 1), \tag{S38a}$$

$$\mu_N > 0. \tag{S38b}$$

Noting that $\lambda^*$ is a decreasing function of $\alpha$ and $\beta$, we find that $\alpha$ and $\beta$ destabilizes the system respectively (Figure S7). The latter result again agrees qualitatively with that of CR and AM, but notably is different from the case when herbivore damage reduces plant fecundity.

### Appendix 3
### Evolution of mean herbivory

#### 6 Introduction

Here, we explore how nonlinearity in the herbivory damage function affects the evolutionary equilibria of mean herbivory. The amount of damage on a plant is a canonical example of a *joint* phenotype *sensu* (34), a trait that is affected by both the evolution of the plant and the herbivore. By analyzing a simple coevolutionary model, we illustrate qualitatively a potential reason why the typical level of damage plants receive in nature tends to be unimodal and not too detrimental to fitness. Importantly, the fact that plants do not routinely experience detrimental levels of herbivore damage is a necessary condition for the herbivory paradox to occur. We begin by briefly summarizing the rationale behind the model structure before showing some key qualitative results.

We examined the evolutionary dynamics of mean herbivore damage $\langle h \rangle$ of a population of plants, a joint phenotype, subject to some coevolutionary dynamics between plant defense and herbivore adaptations to plant defenses (82, 83). To keep the definition of the damage function consistent, we continued to model the dynamics in continuous time as in Appendix 2. Building on (34, 84), the rate of change in the mean phenotype is proportional to the heritable additive genetic variance $G \in (0, \infty]$, which we assume is fixed, and the derivative of the Malthusian fitness function with respect to the mean phenotype $\frac{\partial}{\partial \langle h \rangle} \langle r \rangle (\langle h \rangle)$. To examine the effect of different functional forms of this derivative, we assume that it is fixed and identical across all plant individuals within a population. Ignoring variation in the distribution of damage within population, we can simply replace this derivative with the derivative of our parametric damage function $\frac{\partial}{\partial \langle h \rangle} f(\langle h \rangle_t; \alpha, \beta) = \frac{\beta}{\alpha} \langle h \rangle_t^{\frac{1}{\alpha}-1}$ to analyze its behavior (note that unlike in Appendix 2, the vital rate is subsumed in $\beta$ as is in the main text, but we assume $\beta > 0$ for the sake of discussion). The gradient of the damage function dictates how fast plant defenses will evolve, reducing the herbivore damage that lowers plant fitness.

On the other hand, herbivores also contribute to the evolution of mean herbivore damage. Instead of lowering mean damage, natural selection could favor herbivore adaptions that allow herbivores to eat more plant material. These counter adaptations are known to be widespread (83). Because we are not interested in its influence *per se*, the simplest way we can model the effect of herbivore is with a constant $\nu \in (0, \infty]$, whose strength declines linearly with mean damage. The results remain qualitatively the same even if we set the effect of herbivore to a constant, dropping the $1 - \langle h \rangle$ term. We included the term to stop herbivores from evolving over-exploitative strategies, effectively preventing the evolution of total plant consumption on average, a non-physical state-space. With these ingredients, we can write the coevolutionary dynamics $\kappa(\langle h \rangle_t, t)$ as,

$$\kappa(\langle h \rangle_t, t) = \underbrace{-G \, \frac{\partial}{\partial \langle h \rangle} f(\langle h \rangle_t; \alpha, \beta)}_{\text{Plant defense}} + \underbrace{\nu(1 - \langle h \rangle_t)}_{\text{Herbivore adaptation}} \quad . \tag{S39}$$

The choices of the specific values for the parameters $G$ and $\nu$ do not matter, as they can be scaled relative to $\beta$ to obtain the same qualitative result. That is, what matters is the relative rate of evolution between plants and herbivores. In contrast to other models of coevolution, the implicit treatment of herbivore and plant density implies that both plant and herbivore populations are large, can reproduce regardless of the value of $\langle h \rangle$, and their interactions are diffuse. Similar plant invulnerability at total consumption has been built into other models of plant-herbivore coevolution (85) as is the assumption that herbivores have other food sources to sustain the population when they cannot feed on the focal plant species (86).

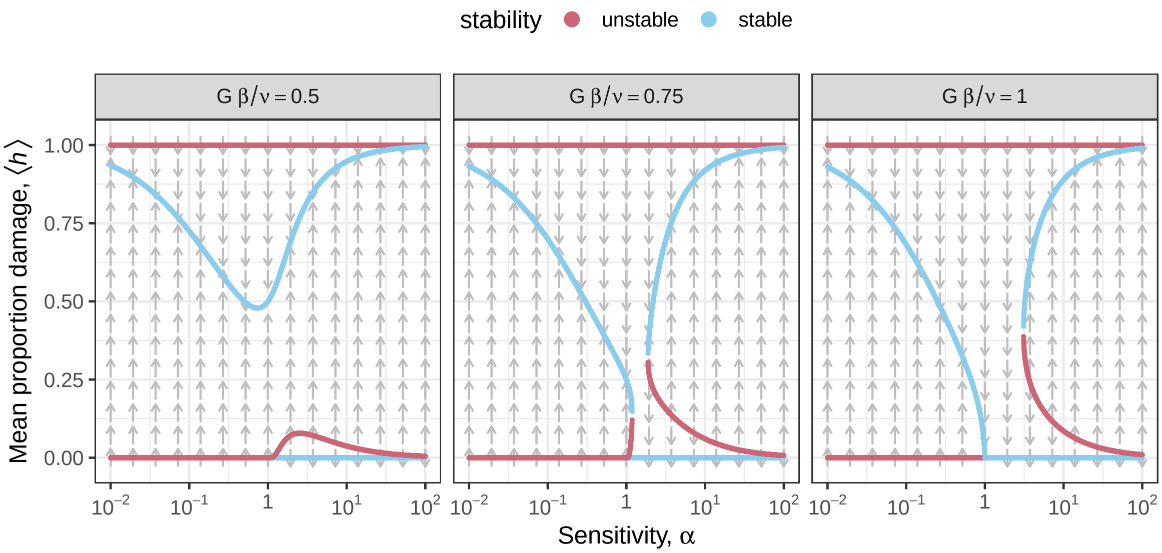

**Figure S8: Equilibria of the simple stochastic mean proportion damage evolution model across different levels of damage sensitivity $\alpha$.** Points show equilibria colored by stability. Arrows show the mean trajectory of the evolution. The system can exhibit bistability when $\alpha > 1$. Numeric solutions and stability were computed using (package rootSolve *87*).

To make the model just slightly more realistic (for a reason that will become clear in the next section), we further assume that the evolutionary dynamics of mean herbivore damage evolve according to Brownian motion with rate $\sigma \in [0, \infty]$ and that evolution beyond the support of proportion damage $h \in [0, 1]$ is clipped at the boundary. Accordingly, we can write a stochastic differential equation of the system as,

$$d\langle h\rangle_t = \underbrace{\kappa(\langle h\rangle_t, t)dt}_{\text{Deterministic coevolution}} + \underbrace{\sigma dW_t}_{\text{Brownian forcing}} + \underbrace{dL_t^0 + dL_t^1}_{\text{Boundary clipping}}. \tag{S40}$$

### 7 Model analysis

The solutions to (S40) cannot be expressed in closed form, but their behavior is fairly straightforward (Figure S8). The key point of this model is to illustrate that plants sensitive to low levels of damage, with $\alpha > 1$, can exhibit bistable evolution of mean herbivore damage, one at 0 and another at a fairly high level. Indeed, without solving for (S40), we can see that the number of solutions depends on $\alpha$. Referring to the first and second terms of (S39) more concisely as $c_1(\langle h\rangle_t)$ and $c_2(\langle h\rangle_t)$, we have solutions at the intersection(s) between $c_1(\langle h\rangle_t)$ and $-c_2(\langle h\rangle_t)$. From (S39), we know that $-c_2(\langle h\rangle_t)$ is a positive line and can only intersect $c_1(\langle h\rangle_t)$ more than once if $c_1(\langle h\rangle_t)$ is concave, which only occurs when $\alpha > 1$.

Stated another way, bistability occurs because if the damage starts out low, the magnitude of selection gradient is very high, so evolution tends towards zero (Figure S9A-B). In contrast, when the damage starts out high, the selection gradient is close to 0, so the herbivore can easily out-evolve the plant defenses. As sensitivity increases, the lower stable equilibrium becomes increasingly untenable, as small stochastic fluctuations can allow the system to jump to the alternative stable state (Figure S9C). This result is important because the evolution of all or nothing levels of damage can lead to frequent catastrophic effects of herbivory on plants, a pattern (bimodal distribution) that is rarely observed in nature (*65*). On the other hand, insensitivity leads to a single stable level of damage that is not as damaging to plant fitness (Figure S9A,C). As a result, it produces the pattern that a large reduction in plant fitness due to herbivore damage is rare (Figure S9D). Of course, for the reasonable range of sensitivities in the neighborhood of $\alpha \sim 1$, these nontrivial qualitative results occur only when the rate of plant and herbivore evolution are on par, $G\beta \sim \nu$. Otherwise, the system fluctuates near a single stable equilibrium at $\langle h \rangle \simeq 1$ or $\langle h \rangle = 0$ depending on whether the plant or herbivore evolves more rapidly.

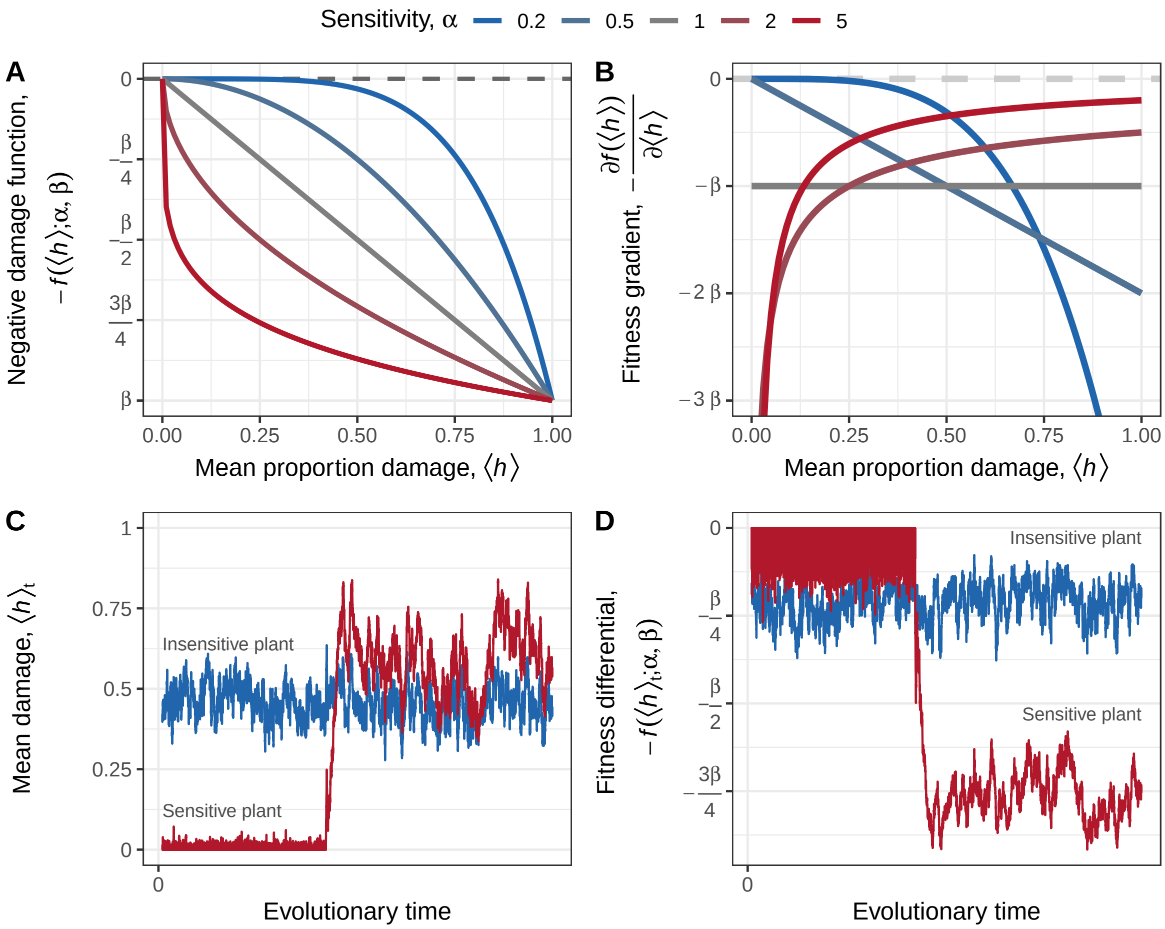

**Figure S9: Evolutionary dynamics of different damage functions.** (A) The fitness differential between a population of plants receiving $\langle h \rangle$ amount of damage compared to a population of plants receiving no damage. Lines are colored by the degree of sensitivity to low levels of herbivore damage $\alpha$. (B) The fitness gradients of the same functions in (A) show that the gradients of sensitive functions tends to decrease with average proportion damage $\langle h \rangle$ and vise versa. (C) The fact that selection gradient of plants sensitive to low levels of damage decreases with increasing proportion damage results in bistability as illustrated in this stochastic simulation with $\sigma = 0.12$, $\beta = 0.6$, $\nu = 1$, and $\alpha = 0.5$ or 2. (D) The same evolutionary trajectories of mean proportion damage $\langle h \rangle_t$ in (C) projected as fitness differential reveal that plants which are insensitive to low levels of damage experience relatively low fitness reduction due to herbivore damage.

### Appendix 4
### Supplemental results

#### 8 Robustness checks

##### 8.1 Appropriateness of damage function

###### 8.1.1 Bitonicity

In this section, we addressed the question of whether the monotonic response (the effect of damage is strictly positive or negative) assumed by our damage function $f(h;\tilde{\alpha},\tilde{\beta})$ adequately describes the empirical data. A long line of research proposes the hypothesis that herbivore damage induces hormetic plant responses, where low levels of damage enhance plant fitness, but high levels of damage reduce plant fitness (*2*, *35*). Given that our damage function is strictly monotonic, such a hormetic response, or any bitonic responses (where the low-damage response is in the opposite direction as the high-damage response) more generally cannot be captured by the model. Therefore, to assess the robustness of our results, we sought to quantify the prevalence of bitonic responses in our dataset.

To do so, we modified the damage function in (S2) to accommodate bitonic responses by introducing a linear term with an opposing sign (Figure S10),

$$\tilde{u}(h;\tilde{\alpha},\tilde{\beta},\tilde{\gamma},\tilde{C}) \equiv \tilde{C} - \tilde{\beta}h^{1/\tilde{\alpha}} + \tilde{\beta}\tilde{\gamma}h. \tag{S41}$$

The inclusion of the leading coefficient $\tilde{\beta}$ scales the bitonic intensity coefficient $\tilde{\gamma}$ relative to $\tilde{\beta}$, so that $\tilde{\gamma}$ can be comparable across datasets and gains an intuitive interpretation of magnitude. If $\tilde{\gamma}$ is too small, then $\tilde{u}(h;\tilde{\alpha},\tilde{\beta},\tilde{\gamma},\tilde{C})$ reduces approximately to $\tilde{C} - \tilde{f}(h;\tilde{\alpha},\tilde{\beta})$. Hence, we can define when the monotonic and bitonic functions are approximately equivalent by defining a Region of Practical Equivalence (ROPE) for $\tilde{\gamma}$. We say that a bitonic response is too small to matter when it is less than 5% of the magnitude of $\tilde{\beta}$, or in other words, when $\ln(\tilde{\gamma}) < -3$. We estimated (S41) using the same procedures as above. We included an additional prior of $\ln(\tilde{\gamma}) \sim \mathcal{N}(-5, 3)$. Using the method described in section 3.2, we can estimate the proportion of datasets that support a bitonic response, assuming a null hypothesis point mass at $\ln(\tilde{\gamma}) = -3$ by shifting $\ln(\tilde{\gamma})$ to the right by 3.

We found that 9.1% of our datasets showed support for a bitonic response and only 5.3% displayed a bitonic response in which plant fitness increased at low levels of damage and decreased at high levels of damage. Therefore, we conclude that our monotonic assumption built into our parametric damage function is an appropriate approximation.

###### 8.1.2 Sigmoid function

In this section, we address the question of whether the monophasic response (having only a monotonic derivative) assumed by our damage function $f(h;\tilde{\alpha},\tilde{\beta})$ adequately describes the empirical data. Some workers propose the hypothesis that the damage function should look sigmoidal, where plants can tolerate low levels of damage and the effect of high levels of damage saturates (*14*, *88*). Given that our damage function is strictly monophasic, such a biphasic sigmoidal response cannot be captured. Therefore, to assess the robustness of our results, we sought to quantify the prevalence of sigmoidal responses in our dataset.

Our analytical strategy follows closely from the previous section. We can generalize the damage function to include both a sigmoidal function and the assumed power function (S8) as special cases. We introduce the term $\tilde{\zeta}$, which controls the degree of sigmoidal shape when $\tilde{\alpha} < 1$, writing (Figure S10),

$$\tilde{v}(h;\tilde{\alpha},\tilde{\beta},\tilde{\zeta},\tilde{C}) \equiv \tilde{C} - \frac{\tilde{\beta}(1+\tilde{\zeta})h^{1/\tilde{\alpha}}}{1+\tilde{\zeta}h^{1/\tilde{\alpha}}}. \tag{S42}$$

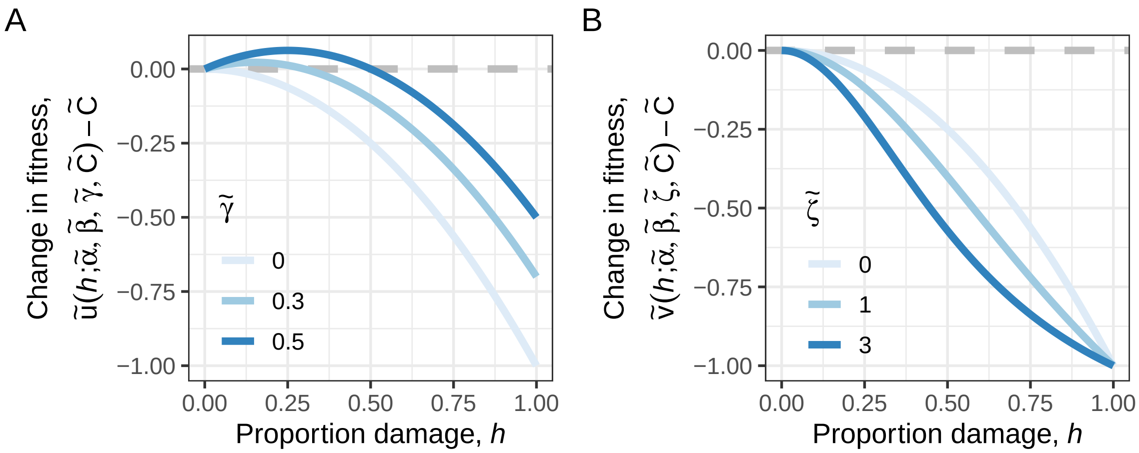

**Figure S10: Generalized damage function families.** (A) A generalized family of functions (S41) that allows for overcompensation. (B) A generalized family of functions (S42) that allows for sigmoid response. In both cases, $\tilde{\beta} = 1$ and $\tilde{\alpha} = 0.5$.

Clearly, when $\tilde{\zeta} = 0$, $\tilde{v}(h; \tilde{\alpha}, \tilde{\beta}, \tilde{\zeta}, \tilde{C})$ reduces to $\tilde{C} - \tilde{f}(h; \tilde{\alpha}, \tilde{\beta})$. Again, we scaled $\tilde{\zeta}$ relative to $\tilde{\beta}$ so that it is comparable across datasets. As before, we say that $\tilde{\zeta}$ is too small to matter when $\ln(\tilde{\zeta}) < 0$, which can be used to classify damage functions. We estimated (S42) using the same procedures as above, with an an additional prior of $\ln(\tilde{\zeta}) \sim \mathcal{N}(-5, 3)$ and assumed a null hypothesis point mass at $\ln(\tilde{\zeta}) = 0$.

We found that only 3.6% of our datasets show support for a sigmoidal response and only 3.1% showed the specific kind of sigmoidal response hypothesized by workers (*14*, *88*) where plant fitness decreased as damage increased. We therefore conclude that our monophasic assumption built into our parametric damage function is an appropriate approximation.

##### 8.1.3 Nonparametric approach

While the parametric form provides interpretable parameters, its structure can be quite restrictive. Therefore, we also fitted more flexible nonparametric functions to check if our analysis is robust to the assumption of the functional form of $f(h)$. By using generalized additive models (GAMs), we are also more able to correctly model the conditional distribution of the data. We fitted the following model using generalized cross-validation, which offers good speed performance (package MGCV *89*),

$$\tilde{Y}|h \sim P(\langle \tilde{Y}|h \rangle, \vartheta), \tag{S43a}$$

$$g(\langle \tilde{Y}|h \rangle) = \tilde{a}_0 - \tilde{f}(h). \tag{S43b}$$

Here, $P(\langle \tilde{Y}|h \rangle, \vartheta)$ denotes the conditional distribution, with the axillary family-specific parameter $\vartheta$; $g(\cdot)$ denotes the link function, $\tilde{a}_0$ denotes the intercept, and $\tilde{f}(h)$ denotes the estimated nonparametric damage function of interest. More precisely, $-\tilde{f}(h)$ is a thin-plate smoother with a basis complexity of four (*90*), which we found in simulations to balance under- and over-fitting. The choice of the link function depended on the assumed distribution of $\tilde{Y}|h$. For beta and gamma distributions, we set $g(x) \equiv \ln(x)$. For the normal distribution, we assumed the variable is already on a log-scale (and for those variables that are log-normally distributed, we log transformed them before fitting), so we set $g(x) \equiv x$. The choice of these link functions differed slightly from what is usually used, but follows from the episodes of selection framework. Further, these link functions allowed us to estimate $\tilde{f}(h)$ in a way that ensures it defaults to a linear form if there is no support for a nonlinear function.

Comparison of results from parametric and nonparametric damage function analysis are shown in Figure S11. As one can see, the results generally show a high degree of concordance. Because splines often perform poorly with extrapolations, we did not compute the effects from spline models when the desired location at which to compute them falls outside of the range of observed data.

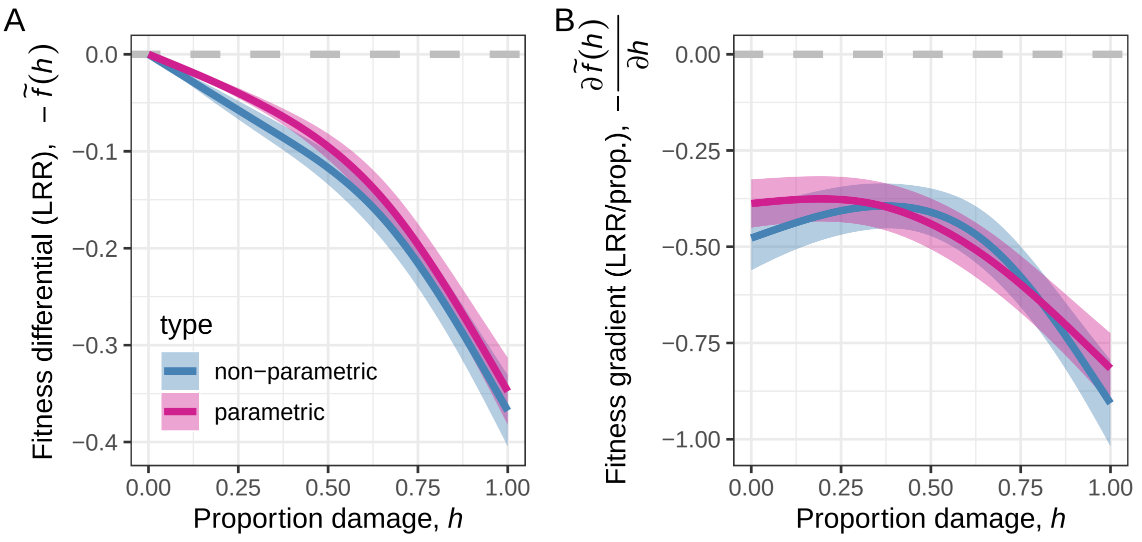

**Figure S11: Comparison of parametric and nonparametric estimates of damage function.** Estimates of (A) average fitness differential and (B) average fitness gradient are similar between effect-sizes derived from splines (S43) and power-functions (S8, S2). Spline models do not assume a family of functional forms, but do not provide effect estimates outside of the range of data. Power-function models do allow for extrapolation, but have a restrictive functional form. Lines and ribbons show mean and 95% confidence intervals. LRR stands for log-response ratio.

#### 8.2 Sensitivity to additional studies

Because the estimated parameters and effects varied significantly across studies, we sought to quantify the robustness of our results to additional studies that may be published in the future. To do so, we calculated a Rosenthal's fail-safe-like number that indicates how many additional studies with an average effect of zero would be required for the estimated effect to become non-significant. We replaced the average $z$-score in Rosenthal's original formulation (eqn. 1 of (*91*)) with the $z$-score of a specific contrast from a fitted multilevel model to accommodate the multilevel structure of our data,

$$k_R = \frac{k\left(z^2 - z_c^2\right)}{z_c^2}. \tag{S44}$$

Here, $k$ is the number of publications in a subgroup and $z_c$ is the critical value at a specific significance level ($z_c = 1.96$ in this study).

#### 8.3 Sources of bias

To explore how the sampling designs of the original studies were related to the parameters we estimated from those studies' data, we fitted two additional multilevel models on $\tilde{\alpha}$ and $\tilde{\beta}$. We used the same model structure as (S10), but dropped the phylogenetic random-effects to improve computational efficiency. For fixed effects, we included log number of data points in the original dataset, log number of unique proportion damage levels measured, and the log range of proportion damage values that was tested. We allowed these fixed effects to vary between damage classes (damaging/benefiting) in the model for $\tilde{\alpha}$.

Results shown in Figure S12 indicate that studies which tested a narrower range of damage levels tended to report more sensitive and less intense effect of simulated herbivore damage. Importantly, this means that our estimation of average fitness differential and fitness gradient may be weakly to moderately positively biased, especially at high-damage levels, if those missing effects were not included in the average. Therefore, in our analysis of fitness differential and fitness gradient, we allowed fitted parametric functions to extrapolate beyond the interval of the observed data. Nevertheless, the results with extrapolation remain fairly similar to results without extrapolation (Figure S11).

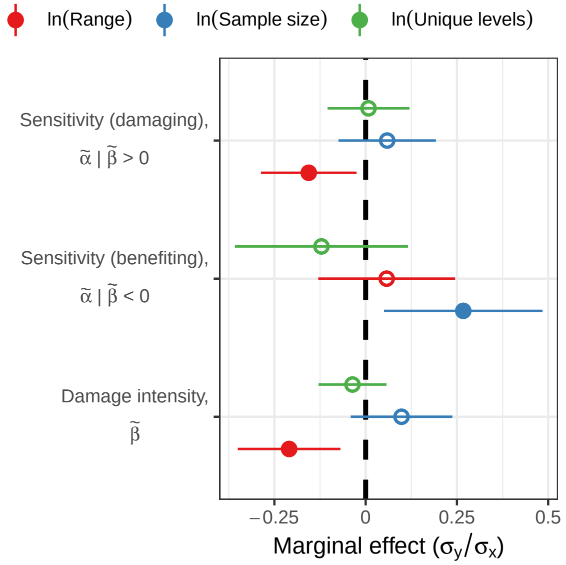

**Figure S12: Marginal effect of sample size, number of unique proportion damage levels, and range of proportion damage tested in the original dataset on the value of damage function parameters.** Points and error bars show mean and 95% confidence intervals. Effects that are significantly different from 0 (black dashed line) or otherwise are displayed as closed and open circles respectively. Because most studies include 0% damage as a control, the range of damage is also approximately the maximum level of damage tested.

### 9 Nonlinear averaging and the herbivory paradox

Here, we explain how nonlinear averaging explains the idea that frequent, low-level herbivore damage are not detrimental to plants and that infrequent, high level herbivore damage maintain the detrimental effects of herbivory on average (*6*, *27*, *28*). Suppose the damage that a plant receives each year is a random variable $H$, if the damage function is accelerating ($\alpha < 1$), then by Jensen's inequality, the average of the damage effects over time must be greater or equal to the damage effects at average damage, $\langle f(H)\rangle \geq f(\langle H\rangle)$. Typical defoliation experiments measure $f(\langle H\rangle)$ instead of $\langle f(H)\rangle$ or $\ln(\langle \exp(f(H))\rangle)$ because the same degree of defoliation is applied to the same plant over time or to all plants in a population.

To estimate the extent that this inequality can inflate the effect of herbivore damage, we need an appropriate distribution of $H$ and an approximation of $f(h)$. As discussed in section 2.1, we can approximate the shape of $f(h)$ with $\tilde{f}(h)$. However, there is insufficient long-term census of $H$ across many species to characterize its distribution well. Fortunately, there are good data on its distribution across space (*29*) and theory suggests that much of this variation may be intrinsic to the stochastic processes of herbivory itself (*30*). Accordingly, the processes that govern the variation in herbivore damage across space may be similar to those across time, so space for time substitution may be a reasonable assumption. We therefore used the individual level damage in the dataset from (*29*) as a stand in for $H$, the typical distribution of herbivore damage. It exhibits properties that are thought to be typical of variation in herbivore damage across time– frequent low-levels of damage and infrequent high levels of damage. Nevertheless, our analysis should be considered qualitative.

By comparing $\langle \tilde{f}(H)\rangle$ with $\tilde{f}(\langle H\rangle)$ for all datasets we analyzed, we can see how nonlinear averaging inflates the magnitude of the effect of herbivore damage (Figure S13A-D). As one expects, with nonlinear averaging, insensitivity to low levels of damage ($\tilde{\alpha} < 1$) inflates the magnitude of the damage effect, whereas sensitivity ($\tilde{\alpha} > 1$) deflates it. But more important is the qualitative behavior that the inflationary effect tends to be much stronger for those effects that were weak to begin with (Figure S13A-B). In effect, nonlinear averaging sets a lower bound to the effect of herbivore damage on a fitness component, and the generally sizable fitness effect at total damage $\tilde{\beta}$ scales the lower bound to be a nontrivial magnitude. This lower bound can also be interpreted as the lower bound to the magnitude of the effect on overall fitness. With nonlinear averaging, the median magnitude changes

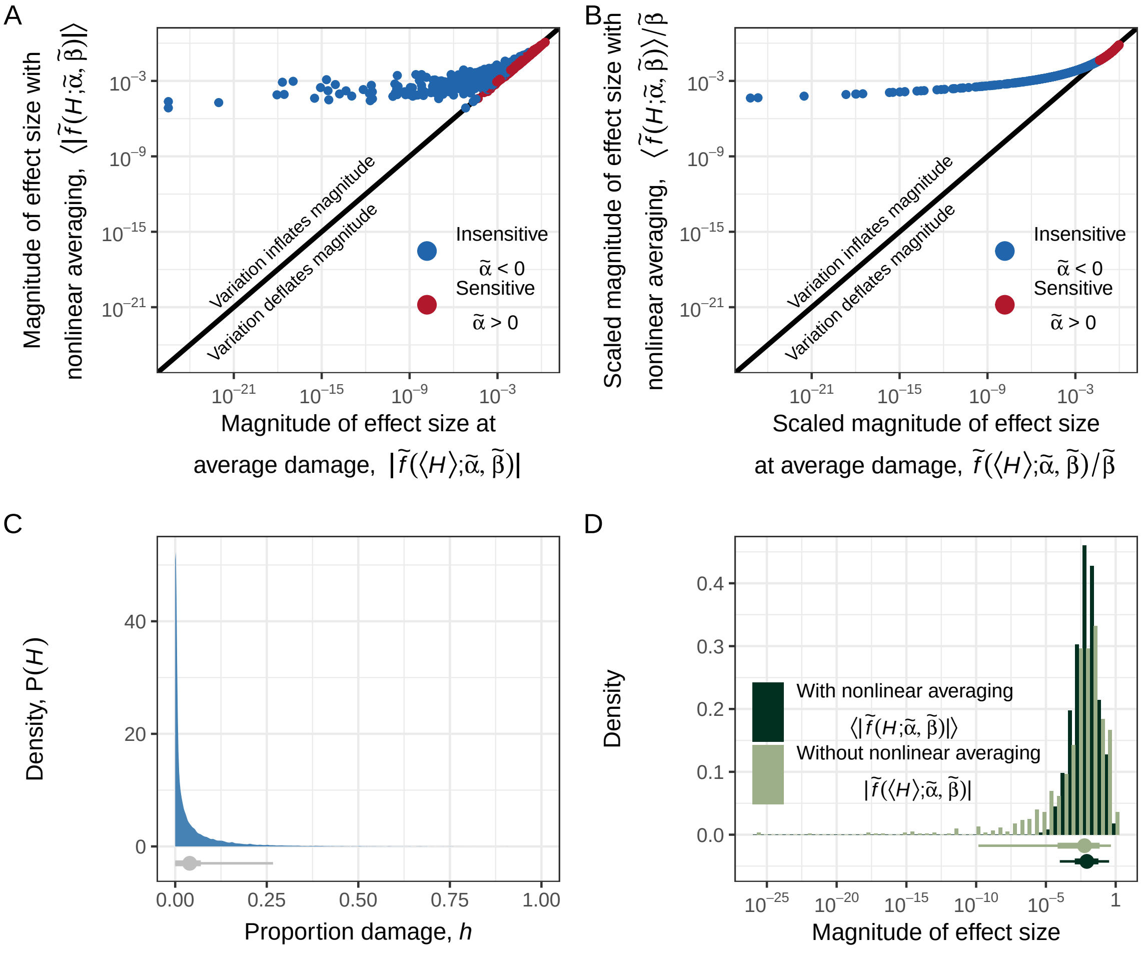

**Figure S13: Damage function nonlinearity and variation in damage prevent the magnitude of damage effect from becoming too small.** (A) The magnitude of damage effects in units of LRR (log-response ratio) with or without nonlinear averaging for a typical distribution of herbivore damage $H$. Each point is a dataset included in our meta-analysis. Points above the identity line indicate that nonlinear averaging inflates the magnitude of the effect, whereas points below the identity line indicate that nonlinear averaging deflates the magnitude. Those effects that are weak to begin with gain the biggest boost in magnitude from nonlinear averaging, effectively setting a lower bound to the importance of herbivore damage. The effects on a fitness component represent lower bounds to the effects on overall fitness. If there are on average a hundred episodes of selection, an effect size of $10^{-4}$ can be relevant to evolution for a population with an effective population size above 1,000. The transition between inflation or deflation of the magnitude by nonlinear averaging is governed by $\tilde{\alpha} = 1$. (B) This transition is most visible when the effects are scaled by $\tilde{\beta}$. (C) The distribution of herbivore damage across time $H$ has a low mean and high skew. The gray point and bars show the 50%, 66%, and 95% quantiles. (D) The distribution of the effect sizes in (A) as histograms. The points and bars show the 50%, 66%, and 95% quantiles.

plant fitness in an episode of selection by 0.86%; without nonlinear averaging, it changes by 0.59%. We should point out that nonlinear averaging does not necessarily make the effect of herbivore damage more important on average across datasets; in fact, the mild deflation of the magnitude of effects that are already strong from sensitive damage functions means that nonlinear averaging leads to a 1.4% increase in plant fitness in an episode of selection on average across datasets. Rather, the key insight is that nonlinear averaging prevents the effect of herbivore damage from becoming not important. A solution to the herbivory paradox suffices that trivial effects of herbivory are generally rendered not trivial. In cases where nonlinear averaging deflates the effect of herbivore damage, these effects are evidently already strong (Figure S13A).

### Appendix 5
### Extended data

#### 10 Figure notes and supporting information

In this section, we present additional information that accompanies figures presented in the main text.

##### 10.1 Figure 1

In (A), we used $\alpha \in \{0.2, 1, 5\}$ for insensitive, neutral, and sensitive functions respectively. In (B), we used the same $\alpha$ values as in (A) and set $\delta = 0.01, \beta = 0.999, b = 0.1, \mu_V = 0.08, \mu_N = 0.05$, and $c = 100$. We simulated the population trajectories for 100 time steps using the reduction in fecundity model (S23). In (C), we used $\alpha \in \{0.5, 2\}$ for insensitive and sensitive functions respective. We set $\beta = 0.6, \nu = 1, \sigma = 0.12$ and simulated the trajectories for 100 time steps.

##### 10.2 Figure 2

**Table S6: Estimates of the average fitness differential and fitness gradient along proportion damage** $h$. Significant parameters are bolded. Estimates show mean [95% confidence intervals] (differential: log-response ratio; gradient: log-response ratio per proportion damage). The locations along proportion damage displayed correspond to where effects from individual damage functions were computed and used for interpolation. The number of effects ($n$), studies (studies), species (species), and a Rosenthal's fail-safe-like number (fail safe) are shown next to each subgroup. Heterogeneity in the true effect, as opposed to sampling error, accounts for most of the variance in fitness differential and fitness gradient ($I^2 > 0.99$).

| Variable | $h$ | Estimate | n | Studies | Species | Fail safe |
|---|---|---|---|---|---|---|
| Fitness differential | 0.05 | **-0.0079 [-0.0092, -0.0066]** | 1112 | 98 | 102 | 3514 |
| | 0.15 | **-0.024 [-0.028, -0.021]** | 1090 | 97 | 100 | 4090 |
| | 0.25 | **-0.043 [-0.049, -0.037]** | 1095 | 98 | 101 | 4814 |
| | 0.5 | **-0.11 [-0.12, -0.093]** | 1049 | 95 | 97 | 4924 |
| | 0.75 | **-0.21 [-0.23, -0.19]** | 877 | 79 | 85 | 7085 |
| | 0.95 | **-0.32 [-0.35, -0.29]** | 619 | 52 | 63 | 5449 |
| Fitness gradient | 0.05 | **-0.38 [-0.44, -0.32]** | 1112 | 98 | 102 | 3713 |
| | 0.15 | **-0.37 [-0.43, -0.31]** | 1090 | 97 | 100 | 3722 |
| | 0.25 | **-0.37 [-0.43, -0.32]** | 1095 | 98 | 101 | 3805 |
| | 0.5 | **-0.46 [-0.53, -0.39]** | 1049 | 95 | 97 | 4374 |
| | 0.75 | **-0.62 [-0.69, -0.54]** | 877 | 79 | 85 | 5330 |
| | 0.95 | **-0.77 [-0.85, -0.68]** | 619 | 52 | 63 | 4092 |

##### 10.3 Figure 3

**Table S7: Marginal effects and predictive performance of site-level variables.** Estimates were derived from the same dataset of 864 effect sizes from 68 studies. Significant effects are bolded. Estimate show mean [95% confidence intervals] in units of $\sigma_y/\sigma_x$. Models are competed against each other for $\tilde{\beta}$, $\tilde{\alpha}|\tilde{\beta} > 0$, and $\tilde{\alpha}|\tilde{\beta} < 0$. An overall metric of fit across all parameters is to sum up the $\text{AIC}_c$ values. Doing so reveals that net primary productivity is the single best predictor ($\Delta\text{AIC}_c$ = 5.3). Heterogeneity in the true effect, as opposed to sampling error, accounts for most of the variance in the damage function parameters for all models ($I^2 > 0.95$).

| Parameter | Variable | Life history | Estimate | $\text{AIC}_c$ | $\Delta\text{AIC}_c$ |
|---|---|---|---|---|---|
| $\tilde{\alpha}\|\tilde{\beta} < 0$ | Absolute latitude | Growth | 0.24 [-0.092, 0.57] | 391.25 | 2.0 |
| | | Reproduction | 0.034 [-0.40, 0.47] | | |
| | | Survival | 0.091 [-0.36, 0.54] | | |
| | Herbivory risk | Growth | -0.013 [-0.33, 0.30] | 392.45 | 3.2 |
| | | Reproduction | -0.26 [-0.74, 0.21] | | |
| | | Survival | -0.58 [-2.7, 1.6] | | |
| | Precipitation | Growth | -0.034 [-0.41, 0.34] | 391.29 | 2.0 |
| | | Reproduction | 0.17 [-0.26, 0.60] | | |
| | | Survival | -0.14 [-0.49, 0.22] | | |
| | Primary productivity | Growth | -0.11 [-0.37, 0.14] | 389.84 | 0.60 |
| | | Reproduction | **0.44 [0.028, 0.85]** | | |
| | | Survival | -0.31 [-0.96, 0.34] | | |
| | Soil fertility | Growth | -0.060 [-0.42, 0.30] | 392.35 | 3.1 |
| | | Reproduction | **-0.37 [-0.69, -0.044]** | | |
| | | Survival | -0.041 [-0.51, 0.43] | | |
| | Temperature | Growth | -0.14 [-0.48, 0.21] | 389.24 | 0.00 |
| | | Reproduction | -0.43 [-0.88, 0.019] | | |
| | | Survival | -0.068 [-0.55, 0.42] | | |
| $\tilde{\alpha}\|\tilde{\beta} > 0$ | Absolute latitude | Growth | -0.074 [-0.22, 0.071] | 1894.11 | 1.6 |
| | | Reproduction | **-0.27 [-0.45, -0.096]** | | |
| | | Survival | 0.17 [-0.45, 0.78] | | |
| | Herbivory risk | Growth | 0.014 [-0.12, 0.15] | 1902 | 9.5 |
| | | Reproduction | -0.023 [-0.20, 0.15] | | |
| | | Survival | -0.30 [-0.79, 0.18] | | |
| | Precipitation | Growth | 0.051 [-0.086, 0.19] | 1892.51 | 0.00 |
| | | Reproduction | **0.26 [0.11, 0.41]** | | |
| | | Survival | -0.038 [-0.41, 0.34] | | |
| | Primary productivity | Growth | -0.11 [-0.24, 0.017] | 1899.19 | 6.7 |
| | | Reproduction | -0.045 [-0.20, 0.11] | | |
| | | Survival | -0.38 [-0.95, 0.19] | | |
| | Soil fertility | Growth | -0.091 [-0.24, 0.061] | 1895.33 | 2.8 |
| | | Reproduction | **-0.23 [-0.40, -0.052]** | | |
| | | Survival | -0.24 [-0.86, 0.39] | | |
| | Temperature | Growth | 0.079 [-0.067, 0.22] | 1897.35 | 4.8 |
| | | Reproduction | **0.22 [0.040, 0.40]** | | |
| | | Survival | -0.094 [-0.75, 0.56] | | |
| $\tilde{\beta}$ | Absolute latitude | Growth | 0.15 [-0.012, 0.32] | 2347.45 | 17 |
| | | Reproduction | **0.22 [0.030, 0.42]** | | |
| | | Survival | **-0.42 [-0.75, -0.091]** | | |
| | Herbivory risk | Growth | -0.019 [-0.17, 0.13] | 2349.73 | 20 |
| | | Reproduction | -0.12 [-0.31, 0.066] | | |
| | | Survival | **0.60 [0.19, 1.0]** | | |
| | Precipitation | Growth | -0.028 [-0.19, 0.13] | 2353.55 | 23 |
| | | Reproduction | -0.16 [-0.34, 0.016] | | |
| | | Survival | 0.24 [-0.0053, 0.49] | | |
| | Primary productivity | Growth | 0.12 [-0.030, 0.27] | 2330.1 | 0.00 |
| | | Reproduction | **-0.25 [-0.43, -0.077]** | | |
| | | Survival | **0.54 [0.16, 0.93]** | | |
| | Soil fertility | Growth | -0.093 [-0.27, 0.080] | 2355.99 | 26 |
| | | Reproduction | -0.053 [-0.24, 0.13] | | |
| | | Survival | **0.34 [0.0023, 0.67]** | | |
| | Temperature | Growth | -0.14 [-0.31, 0.024] | 2337.87 | 7.8 |

| | | | |
|---|---|---|---|
| | | Reproduction | **-0.37 [-0.57, -0.18]** |
| | | Survival | **0.52 [0.16, 0.88]** |

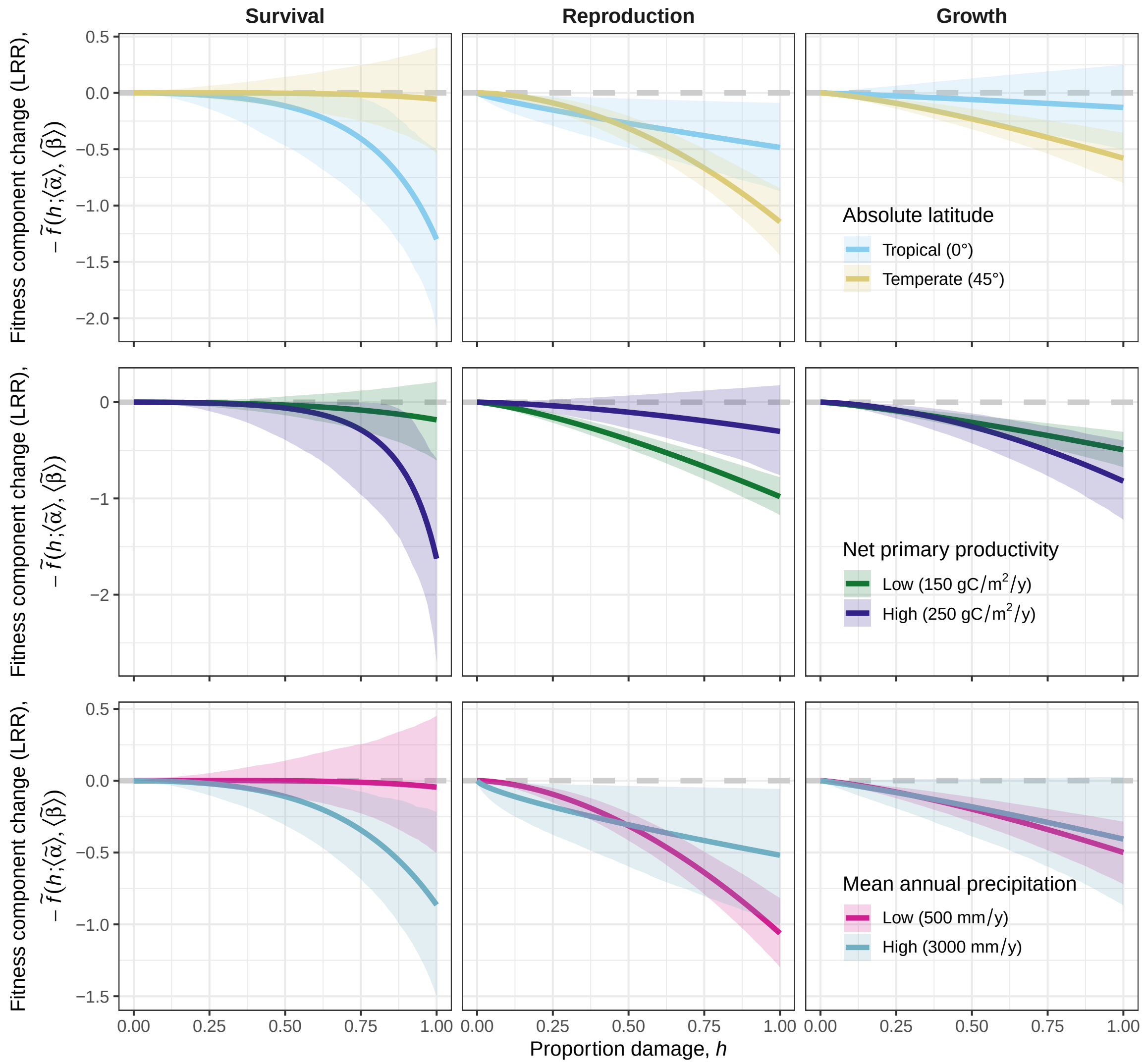

**Figure S14: Negative damage function for different subgroups analyzed in Figure 3 in the main text.** Each curve is drawn using the predicted average parameter for the corresponding group. To account for uncertainty in the estimated damage function parameters, we performed 2,000 bootstraps using the mean and standard error of the parameter estimates. The correlation between $\tilde{\alpha}$ and $\tilde{\beta}$ is assumed to be -0.22 based on trends in the raw data. Lines and ribbons show mean and 95% confidence intervals. LRR stands for log-response ratio. The first row of panels are identical to the ones in the main text.

#### 10.4 Figure 4

**Table S8: Estimates of damage function parameters across subgroups.** Significant parameters are bolded. Estimate show mean [95% confidence intervals] (log-response ratio for $\tilde{\beta}$; unitless for $\tilde{\alpha}$). The number of effects ($n$), studies (studies), species (species), and a Rosenthal's fail-safe-like number (fail safe) are shown next to each subgroup. Damage organ best explains $\tilde{\alpha}$ ($\Delta\text{AIC}_c = 4.9$) and life history best explains $\tilde{\beta}$ ($\Delta\text{AIC}_c = 17.6$) and both parameters jointly ($\Delta\text{AIC}_c = 22.7$). Heterogeneity in the true effect, as opposed to sampling error, accounts for most of the variance in the damage function parameters for all models ($I^2 > 0.96$).

| Variable | Condition | Parameter | Estimate | n | Studies | Species | Fail safe |
|---|---|---|---|---|---|---|---|
| Life history | Growth | $\tilde{\alpha}\|\tilde{\beta} < 0$ | 1.1 [0.87, 1.4] | 113 | 32 | 38 | 0 |
| | | $\tilde{\alpha}\|\tilde{\beta} > 0$ | **0.83 [0.71, 0.96]** | 531 | 73 | 78 | 53 |
| | | $\tilde{\beta}$ | **0.54 [0.39, 0.70]** | 644 | 75 | 84 | 835 |
| | Reproduction | $\tilde{\alpha}\|\tilde{\beta} < 0$ | **1.5 [1.1, 1.9]** | 74 | 29 | 22 | 26 |
| | | $\tilde{\alpha}\|\tilde{\beta} > 0$ | **0.79 [0.67, 0.94]** | 395 | 52 | 40 | 50 |
| | | $\tilde{\beta}$ | **0.86 [0.69, 1.0]** | 469 | 52 | 40 | 1295 |
| | Survival | $\tilde{\alpha}\|\tilde{\beta} < 0$ | 0.67 [0.35, 1.3] | 12 | 5 | 7 | 0 |
| | | $\tilde{\alpha}\|\tilde{\beta} > 0$ | **0.46 [0.30, 0.71]** | 20 | 9 | 9 | 20 |
| | | $\tilde{\beta}$ | 0.20 [-0.13, 0.54] | 32 | 12 | 15 | 0 |
| Organ | Leaf | $\tilde{\alpha}\|\tilde{\beta} < 0$ | 1.2 [1.0, 1.5] | 159 | 45 | 44 | 0 |
| | | $\tilde{\alpha}\|\tilde{\beta} > 0$ | **0.82 [0.71, 0.95]** | 804 | 83 | 73 | 78 |
| | | $\tilde{\beta}$ | **0.58 [0.43, 0.73]** | 963 | 83 | 77 | 1132 |
| | Stem/Shoot | $\tilde{\alpha}\|\tilde{\beta} < 0$ | 0.78 [0.47, 1.3] | 26 | 5 | 11 | 0 |
| | | $\tilde{\alpha}\|\tilde{\beta} > 0$ | **0.59 [0.43, 0.81]** | 102 | 11 | 20 | 21 |
| | | $\tilde{\beta}$ | **0.86 [0.50, 1.2]** | 128 | 12 | 24 | 58 |
| | Root | $\tilde{\alpha}\|\tilde{\beta} < 0$ | **43 [7.4, 250]** | 1 | 1 | 1 | 4 |
| | | $\tilde{\alpha}\|\tilde{\beta} > 0$ | 0.89 [0.23, 3.4] | 2 | 1 | 1 | 0 |
| | | $\tilde{\beta}$ | 0.57 [-0.57, 1.7] | 3 | 1 | 1 | 0 |
| | Flower/Seed/Fruit | $\tilde{\alpha}\|\tilde{\beta} < 0$ | 0.77 [0.35, 1.7] | 13 | 3 | 3 | 0 |
| | | $\tilde{\alpha}\|\tilde{\beta} > 0$ | 1.0 [0.61, 1.8] | 32 | 6 | 6 | 0 |
| | | $\tilde{\beta}$ | **1.0 [0.46, 1.6]** | 45 | 6 | 6 | 14 |
| | Unknown | $\tilde{\alpha}\|\tilde{\beta} < 0$ | - | 0 | 0 | 0 | - |
| | | $\tilde{\alpha}\|\tilde{\beta} > 0$ | 0.56 [0.16, 1.9] | 6 | 1 | 1 | 0 |
| | | $\tilde{\beta}$ | 1.0 [-0.35, 2.4] | 6 | 1 | 1 | 0 |
| Damaged stage | Seed | $\tilde{\alpha}\|\tilde{\beta} < 0$ | 1.3 [0.19, 8.9] | 2 | 1 | 1 | 0 |
| | | $\tilde{\alpha}\|\tilde{\beta} > 0$ | 1.4 [0.56, 3.5] | 13 | 2 | 2 | 0 |
| | | $\tilde{\beta}$ | **1.8 [1.0, 2.6]** | 15 | 3 | 3 | 14 |
| | Seedling | $\tilde{\alpha}\|\tilde{\beta} < 0$ | 0.94 [0.54, 1.6] | 26 | 5 | 11 | 0 |
| | | $\tilde{\alpha}\|\tilde{\beta} > 0$ | 0.80 [0.55, 1.2] | 52 | 10 | 22 | 0 |
| | | $\tilde{\beta}$ | **0.47 [0.12, 0.81]** | 78 | 11 | 27 | 9 |
| | Vegetative | $\tilde{\alpha}\|\tilde{\beta} < 0$ | 1.2 [0.95, 1.6] | 98 | 30 | 31 | 0 |
| | | $\tilde{\alpha}\|\tilde{\beta} > 0$ | **0.78 [0.67, 0.91]** | 544 | 66 | 58 | 105 |
| | | $\tilde{\beta}$ | **0.57 [0.42, 0.72]** | 642 | 67 | 61 | 913 |
| | Flowering | $\tilde{\alpha}\|\tilde{\beta} < 0$ | 1.2 [0.77, 1.8] | 38 | 14 | 14 | 0 |
| | | $\tilde{\alpha}\|\tilde{\beta} > 0$ | **0.79 [0.63, 0.99]** | 171 | 27 | 23 | 2 |
| | | $\tilde{\beta}$ | **0.59 [0.39, 0.79]** | 209 | 28 | 23 | 213 |
| | Fruiting | $\tilde{\alpha}\|\tilde{\beta} < 0$ | 1.1 [0.67, 1.8] | 27 | 10 | 8 | 0 |
| | | $\tilde{\alpha}\|\tilde{\beta} > 0$ | **0.73 [0.56, 0.95]** | 122 | 17 | 12 | 8 |
| | | $\tilde{\beta}$ | **0.84 [0.61, 1.1]** | 149 | 17 | 12 | 205 |
| | Other | $\tilde{\alpha}\|\tilde{\beta} < 0$ | 1.1 [0.51, 2.2] | 8 | 4 | 5 | 0 |
| | | $\tilde{\alpha}\|\tilde{\beta} > 0$ | 0.90 [0.60, 1.4] | 44 | 9 | 11 | 0 |
| | | $\tilde{\beta}$ | **1.0 [0.60, 1.5]** | 52 | 9 | 11 | 43 |
| Perenniality | Annual | $\tilde{\alpha}\|\tilde{\beta} < 0$ | **1.4 [1.1, 2.0]** | 85 | 23 | 15 | 8 |
| | | $\tilde{\alpha}\|\tilde{\beta} > 0$ | 0.82 [0.65, 1.0] | 425 | 38 | 24 | 0 |
| | | $\tilde{\beta}$ | **0.56 [0.34, 0.79]** | 510 | 38 | 24 | 193 |
| | Perennial | $\tilde{\alpha}\|\tilde{\beta} < 0$ | 1.0 [0.79, 1.3] | 114 | 30 | 40 | 0 |
| | | $\tilde{\alpha}\|\tilde{\beta} > 0$ | **0.79 [0.67, 0.94]** | 521 | 61 | 71 | 59 |
| | | $\tilde{\beta}$ | **0.69 [0.51, 0.87]** | 635 | 62 | 79 | 848 |
| Growth form | Graminoid | $\tilde{\alpha}\|\tilde{\beta} < 0$ | 1.5 [0.99, 2.4] | 38 | 10 | 9 | 0 |
| | | $\tilde{\alpha}\|\tilde{\beta} > 0$ | 0.83 [0.60, 1.1] | 197 | 16 | 14 | 0 |
| | | $\tilde{\beta}$ | **0.52 [0.19, 0.85]** | 235 | 16 | 14 | 24 |
| | Herbaceous | $\tilde{\alpha}\|\tilde{\beta} < 0$ | 1.0 [0.79, 1.4] | 106 | 28 | 23 | 0 |
| | | $\tilde{\alpha}\|\tilde{\beta} > 0$ | **0.77 [0.63, 0.94]** | 477 | 46 | 34 | 36 |
| | | $\tilde{\beta}$ | **0.56 [0.36, 0.76]** | 583 | 47 | 35 | 316 |
| | Climber | $\tilde{\alpha}\|\tilde{\beta} < 0$ | 1.6 [0.72, 3.7] | 10 | 4 | 3 | 0 |
| | | $\tilde{\alpha}\|\tilde{\beta} > 0$ | 0.94 [0.58, 1.5] | 52 | 8 | 6 | 0 |

| | | | | | | | |
|---|---|---|---|---|---|---|---|
| | | $\tilde{\beta}$ | **1.1 [0.64, 1.6]** | 62 | 8 | 6 | 34 |
| | Woody | $\tilde{\alpha}\|\tilde{\beta} < 0$ | 1.1 [0.73, 1.5] | 45 | 13 | 20 | 0 |
| | | $\tilde{\alpha}\|\tilde{\beta} > 0$ | **0.79 [0.62, 1.0]** | 220 | 30 | 41 | 2 |
| | | $\tilde{\beta}$ | **0.70 [0.45, 0.95]** | 265 | 31 | 48 | 212 |
| Domestication | Domesticated | $\tilde{\alpha}\|\tilde{\beta} < 0$ | 1.2 [0.96, 1.6] | 112 | 31 | 22 | 0 |
| | | $\tilde{\alpha}\|\tilde{\beta} > 0$ | **0.73 [0.61, 0.87]** | 553 | 54 | 34 | 113 |
| | | $\tilde{\beta}$ | **0.66 [0.47, 0.85]** | 665 | 54 | 34 | 610 |
| | Wild type | $\tilde{\alpha}\|\tilde{\beta} < 0$ | 1.0 [0.80, 1.4] | 87 | 22 | 33 | 0 |
| | | $\tilde{\alpha}\|\tilde{\beta} > 0$ | 0.88 [0.73, 1.1] | 393 | 47 | 61 | 0 |
| | | $\tilde{\beta}$ | **0.61 [0.42, 0.81]** | 480 | 48 | 69 | 415 |

### 11 Meta-analysis references
